# Unveiling Frequency-Specific Microstate Correlates of Anxiety and Depression Symptoms

**DOI:** 10.1101/2024.03.28.587119

**Authors:** Siyang Xue, Xinke Shen, Dan Zhang, Zhenhua Sang, Qiting Long, Sen Song, Jian Wu

## Abstract

Electroencephalography (EEG) microstates are canonical voltage topographies that reflect the temporal dynamics of resting-state brain networks on a millisecond time scale. Changes in microstate parameters have been described in patients with psychiatric disorders, indicating their potential as clinical biomarkers with broadband EEG signals (e.g., 1–30 Hz). Considering the distinct information provided by specific frequency bands, we hypothesized that microstates in decomposed frequency band could provide a more detailed depiction of the underlying psychological mechanism. In this study, with a large open access resting-state dataset (n = 203), we examined the properties of frequency-specific microstates and their relationship with emotional disorders. We conducted clustering on EEG topographies in decomposed frequency band (delta, theta, alpha and beta), and determined the number of clusters with a meta-criterion. Microstate parameters, including global explained variance (GEV), duration, coverage, occurrence and transition probability, were calculated for eyes-open and eyes-closed states, respectively. Their predictive power for the scores of depression and anxiety symptoms were identified by correlation and regression analysis. Distinct microstate patterns were observed across different frequency bands. Microstate parameters in the alpha band held the best predictive power for emotional symptoms. Microstates B (GEV, coverage) and parieto-central maximum microstate C’ (coverage, occurrence, transitions from B to C’) in the alpha band exhibited significant correlations with depression and anxiety, respectively. Microstate parameters of the alpha band achieved predictive R-square of 0.100 for anxiety scores, which is much higher than those of broadband (R-square = -0.026, p < .01). These results suggested the value of frequency-specific microstates in predicting emotional symptoms.

## 1 Introduction

Currently, research on the brain’s intrinsic activity, or resting-state studies, plays an important role in studies of the human brain in health and disease (Raichle, 2015). Abnormal large-scale functional brain network activities have been indicated in the pathophysiology of emotional disorders, such as depression and anxiety (Anand et al., 2005; Pizzagalli, 2011; Sylvester et al., 2012; Xu et al., 2019). The objective of these studies is to find reliable predictors that can describe differences between healthy individuals and clinical subgroups (Dubois et al., 2018). As a prevalent neuroimaging technique, resting-state functional magnetic resonance imaging (rs-fMRI) is widely used to examine the neurofunctional bases underlying neuropsychiatric disorders (Biswal, 2012; Raichle, 2010; Wang et al., 2019; Feurer et al., 2021).

However, the activity related blood-oxygen-level dependent (BOLD) signal limits the analysis to a time scale of seconds due to hemodynamic response function (Abrol et al., 2017; Preti et al., 2017). Electroencephalography (EEG), with a higher temporal resolution, might thus serve as an effective supplement to fMRI (Bressler, 1995; Martınez-Montes et al., 2004; Michel and Koenig, 2018). This method might be more suitable for describing the rapid modifications that occur within the complex mental activities at rest (Murphy et al., 2020). EEG microstate, which characterizes the quasi-stable properties of EEG activities and reflects the dynamics of large-scale brain networks, has been found to change in a variety of brain disorders (Michel and Koenig, 2018). It provides a potential biomarker for characterizing depression and anxiety symptoms with resting-state EEG.

Microstate (MS) described the dynamics of global electric potentials over multichannel EEG recordings. These topographies are stable for a short time period (∼60-120 ms) and rapidly transition to another quasi-stable topography (Lehmann, 1971; Lehmann et al., 1987). As MS simultaneously considers signals recorded from all cortical regions, it is capable of assessing the function of large-scale brain networks whose disruption is associated with several neuropsychiatric disorders (Van De Ville et al., 2010; Michel and Koenig, 2018), such as schizophrenia (Khanna et al., 2014; Tomescu et al., 2014; Diaz Hernandez et al., 2016; da Cruz et al., 2020; Kim et al., 2021), dementia (Grieder et al., 2016; Lassi et al., 2023) and obsessive-compulsive disorder (Thirioux et al., 2023). Recently, microstate analysis was also applied to depression and anxiety disorders (Al Zoubi et al., 2019; Murphy et al., 2020; Lei et al., 2022; Li et al., 2022; Chivu et al., 2023). These studies revealed discernible differences in EEG microstate parameters between patients with depression and anxiety disorders and general population, with significant effects mainly observed in microstates B and D. However, it should be noted that existing research has predominantly concentrated on contrasting clinical populations with healthy individuals, with scant attention given to the general population exhibiting mild or early-stage symptoms. This study aims to specifically explore the prediction of anxiety and depression symptoms within the general population, filling a notable gap in the existing literature.

Besides, most existing studies only focused on broadband EEG signals, the unique temporal information within frequency-specific activities have been largely neglected. Recent research indicated that, compared to broadband, microstates features in decomposed frequency bands outperformed in distinguishing between eyes-closed and eyes-open states (Férat et al., 2022), and provided a more precise distinction between patients with Post-Traumatic Stress Disorder (PTSD) and the healthy controls (Terpou et al., 2022). Furthermore, these features were also found to be more effective in emotion recognition (Shen et al., 2020), which suggested frequency-specific microstates might reflect the neural substrates of emotion processing better. However, the relationship between frequency-specific microstate and emotional disorders remains underexplored. Given that emotional disorders largely stem from deficits in emotion processing (Etkin and Wager, 2007), an investigation into frequency-specific microstates might provide new insights and hold potentials to predict symptoms of emotional disorders.

To address the above questions, the primary aim of this study was to explore the spatiotemporal dynamics of narrowband microstates during resting-state, further unveiling their associations with symptoms of emotional disorders. To achieve this, we analyzed both eyes-closed (EC) and eyes-open (EO) resting-state EEG (rs-EEG) recordings from LEMON dataset (Babayan et al., 2019). EEG was filtered into broadband (1-30Hz), delta band (1-4 Hz), theta band (4-8 Hz), alpha band (8-12 Hz) and beta band (15-30 Hz) before microstate analysis. The number of clusters for each frequency band was individually determined via the meta-criterion method (Bréchet et al., 2019; Férat et al., 2022). Then we examined the microstate topographies and temporal properties in each frequency band. Topographic Electrophysiological State Source-Imaging (TESS) method was deployed to explore the neural generators of microstates in specific frequency bands (Custo et al., 2017). Further, the relationship between microstate parameters and anxiety or depression symptoms was identified by bivariate partial correlation, controlling for age group differences. Finally, regression models were employed to predict the severity of anxiety or depression symptoms by microstate parameters, comparing those in broadband and in specific frequency bands. We hypothesized that in resting-state, microstate parameters in a specific frequency band exhibit: (i) unique topographic configurations and spatiotemporal dynamics, (ii) significant correlations with emotional disorders, and (iii) improved ability to predict the emotional disorders than those in broadband.

## 2 Materials and methods

### 2.1 Participants

EEG recordings from 227 anonymized participants were provided by Leipzig Study for Mind-Body-Emotion Interactions (LEMON). The participants were divided into two groups: younger adults between 20 and 35 years old (N = 153, 45 females, mean age = 25.1 years, SD = 3.1) and older adults between 59 and 77 years old (N = 74, 37 females, mean age = 67.6 years, SD = 4.7). This dataset collected a large amount of physiological, psychological, and neuroimaging measurement data. Assessment and data collection details were provided in (Babayan et al., 2019).

Thirteen participants were excluded due to missing event information, errors in sampling rate, or insufficient data quality, leaving 203 participants with useable EEG. EEG recordings consist of 62 channels (ActiCAP, Brain Products GmbH, Germany) placed according to the international 10–10 system arrangement, with one additional electrode below the right eye to record vertical eye movements. The reference electrode was located at position FCz and the ground was located at the sternum.

The LEMON dataset was collected over two days, including a 3.0T MRI (day 1) and a 62-channel rs-EEG experiment (16 minutes, day 2). Each EEG recording was taken from participants in a sound-attenuated chamber, divided into 16 contiguous 1-minute blocks, with 8 minutes of eyes-closed and 8 minutes of eyes-open conditions interleaved, starting with the eyes-closed condition. Immediately following, 18 emotion-related questionnaires were arranged in a randomized sequence on a computer, the whole questionnaire completion took on average 1.5 hours to 2.5 hours with a short break after 45 min. The study protocol was approved by the ethics committee of the University of Leipzig (reference 154/13-ff). Data were obtained in accordance with the Declaration of Helsinki.

### 2.2 Preprocessing

As reported by (Babayan et al., 2019), the preprocessing of LEMON dataset involves the following steps: The raw EEG data was initially downsampled from 2500 to 250 Hz and bandpass filtered between 1 and 45 Hz with an 8th order Butterworth filter. This was followed by the removal of abnormal channels and noisy segments. Next, principal component analysis (PCA) was used to reduce the dimensionality of the data, retaining components that account for 95% of the total data variance. Independent component analysis (ICA) was then used to remove artifacts such as eye movements, blinks, and heartbeats for EC and EO conditions.

We further preprocessed the data with the following operations using Python-MNE (Gramfort et al., 2013): The missing electrodes were interpolated using spherical spline interpolation, thus obtaining 61 usable channels. The data were re-referenced by the average of all channels, and downsampled form 250 Hz to 100 Hz. Finally, EEG recordings were filtered into different frequency bands: broadband (1–30 Hz), delta (1–4 Hz), theta (4–8 Hz), alpha (8–12 Hz) and beta (15–30 Hz). The filter is a two-pass, zero-phase and non-causal bandpass finite-impulse response filter.

### 2.3 Microstate analysis

We conducted microstate analysis with Cartool software (Brunet et al., 2011). EC and EO conditions were analyzed separately here. Firstly, local maxima of the global field power (GFP) were extracted from EEG data, and submitted to a modified k-means clustering analysis. The process involved assessing 50 random initializations of the modified k-means algorithm for each number of centroids, ranging from 1 to 12. The initialization with the highest global explained variance (GEV) was selected for further processing. Subsequently, we obtained 4060 clustering results for each condition & frequency band. These results were randomly resampled into 100 subsets and submitted to a modified k-means clustering analysis to extract 100 optimal centroid sets. The final step was merging these sets and extracting a set of k centroids that best represent the spatiotemporal variance of frequency-specific EEG data under various conditions. This method was used to calculate the optimal number of clusters (k) for each frequency band.

Once the cluster centroids were identified, microstate patterns were designated based on correlation analysis both within and between frequencies. Firstly, we compared the topographies between EC and EO conditions within each frequency band. This was done by calculating the spatial correlations to examine whether a topographic configuration differed under different conditions. Next, we conducted the comparison of topographies between broadband and each narrowband. Topographies with high correlation would be defined as the same microstate (r ≥ .90).

### Back-fitting of MS maps

Each time point from all individual recordings was aligned with the topography that exhibited the highest absolute spatial correlation, independently for each frequency and condition. Time points that had low spatial correlation (r < 0.5) with all topographies were designated as ‘unassigned’. Temporal smoothing (Pascual-Marqui et al., 1995; Brunet et al., 2011) was implemented in the following rule: Segments with a duration shorter than three samples (30 ms) were split into two parts, each of which was allocated to the neighboring segment.

### Temporal parameters

Four classical parameters were derived from each microstate. (i) GEV: defined as the percentage of observed topographic variance explained by each specific global topographic centroid, serves as a unique identifier for each topographic map (Murray et al., 2008). (ii) Mean duration: the average duration (measured in milliseconds) of continuous samples from the EEG time series that are categorized according to a specific microstate configuration. In neurophysiological terms, this duration is believed to represent the average length of time during which a group of neural generators maintains synchronous activity (Khanna et al., 2015). (iii) Time coverage: the average prevalence of each microstate map throughout the entire EEG recording, expressed as a percentage. This is calculated by dividing the number of time points assigned to a specific microstate map by the total number of time points. (iv) Occurrence: quantifies the average instances per second in which a specific microstate emerges.

### Transition probabilities

We investigated first order Markov-chain transition probabilities from the microstate segmentations using Cartool. The probability that a given microstate would transition to another microstate configuration was calculated for each pair of microstate configurations for each individual recording.

### Adjusted mutual information score

Adjusted Mutual Information Score (AMI) (Vinh et al., 2010) measures the shared information between two microstate segmentations. A score close to 1 indicates similar temporal patterns, while a score near 0 indicates independent sequences. The adjustment accounts for potential overlaps that could occur by chance.

### Source localization

Source localization was achieved through dSPM (dynamic statistical parametric mapping) (Dale et al., 1999), which estimated the time course of the current density with 17001 solution points (Michel et al., 2004) via the Brainstorm (Tadel et al., 2011). This time course was then fitted with the temporal regressors through GLM (general linear model). Finally, we performed bootstrapping (Trevor Hastie et al., 2009) to examine the location and amplitude of the generators of each microstate, thus determining EEG based resting-state networks (eRSNs). The methodological schema for TESS is depicted in Fig. S1. The overview for microstate analysis can be found in Fig. 1.

**Fig. 1.**
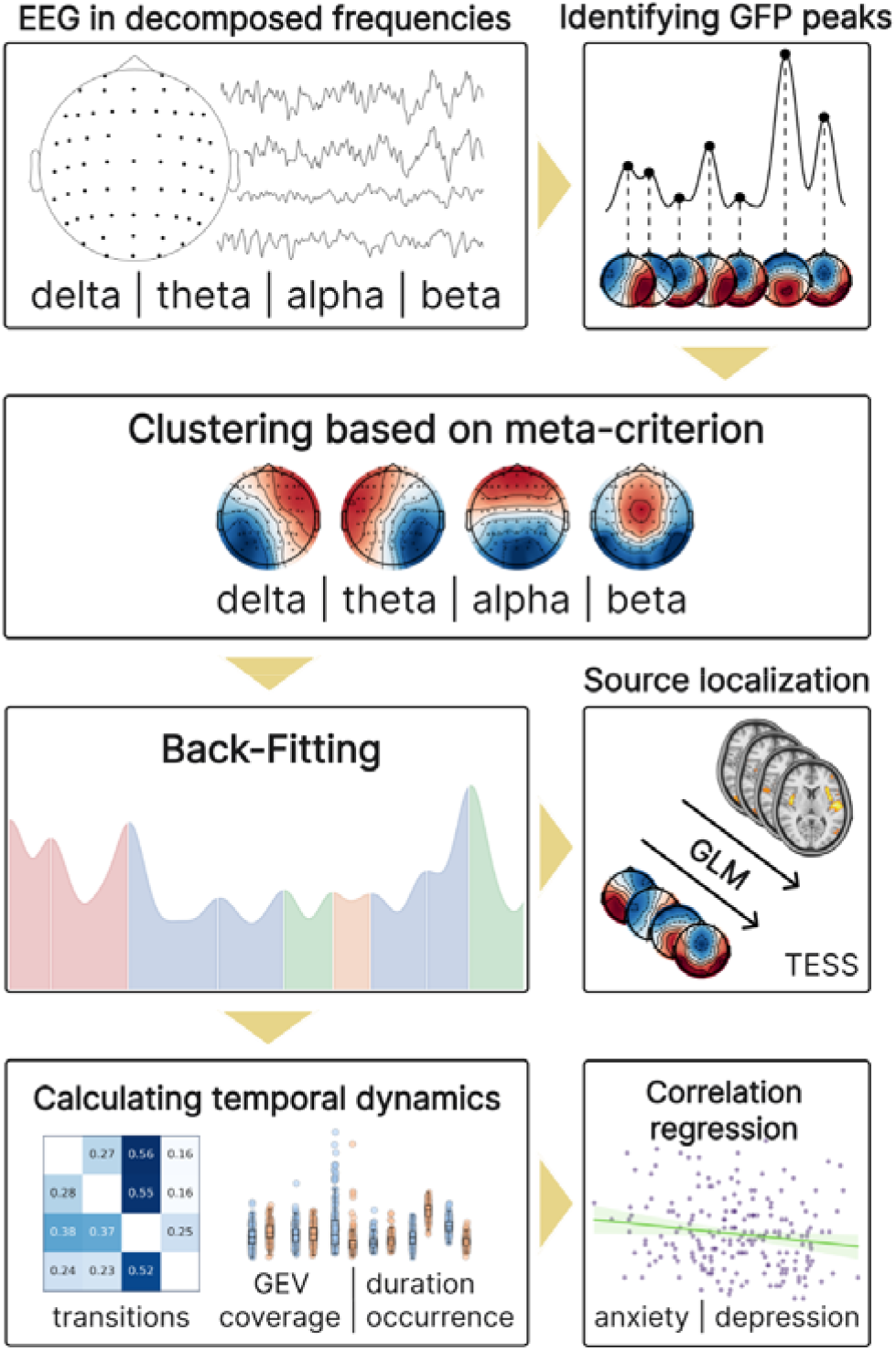
The overview of microstate analysis

### 2.4 Assessment of anxiety and depression symptoms

#### Anxiety

In the State-Trait Anxiety Inventory (STAI-G-X2), trait anxiety (T-Anxiety) is defined in terms of relatively stable individual differences in anxiety-proneness as reflected in the frequency that anxiety states have been manifested in the past and the probability that feelings of anxiety will be experienced in the future (Spielberger et al., 1970). See (Spielberger and Reheiser, 2009) for more detailed interpretation of conceptual framework. T-Anxiety consists of 20 items that exhibit great stability over time and high correlations with the Manifest Anxiety Scale (Taylor, 1953) and Cattell and Anxiety Scale Questionnaire (Krug, 1976). Each item is rated on a 4-point Likert scale ranging from 1 (“almost never”) to 4 (“nearly always”). Individuals with high score of T-Anxiety tend to experience feelings of tension, apprehension, nervousness, and worry more frequently and intensely.

#### Depression

Depressive symptoms are assessed by a trained psychologist or research assistant using the Hamilton Depression Scale (Hamilton, 1960). The evaluation includes the sum score of 17 items, with most items rated on a 5-point Likert scale. The rating standards for each level are defined as: (0) None; (1) Mild; (2) Moderate; (3) Severe; and (4) Extremely Severe.

### 2.5 The contrast of microstate features across frequencies and different states

To examine the differences across frequencies and conditions, we applied paired t-tests to microstate temporal parameters (GEV, duration, coverage and occurrence) and transition probabilities. False Discovery Rate (FDR) (Benjamini and Hochberg, 1995) was applied to p-values for multiple comparisons. We also calculated standardized mean difference (Cohen’s d) (Gibbons et al., 1993) to report the effect sizes.

Random forest classifier was applied to differentiate behavioral states (EC vs. EO). The models were fitted with standardized microstate parameters (GEV, duration, coverage, occurrence and transition probabilities) derived from the given frequency band. The performance was validated with a 10-fold cross-validation. Model performance was evaluated using the mean accuracy (ACC) and area under the curve (AUC). ACC represents the proportion of correctly predicted individuals in the validation set. On the other hand, AUC is defined as the area under the receiver operating characteristic (ROC) curve, considered as an aggregate measure of performance across all possible classification thresholds. Higher values of ACC and AUC indicate better predictive performance.

### 2.6 Correlation and regression analysis between microstate features and anxiety/depression symptoms

For correlation analyses, we performed a bivariate partial correlation between microstate parameters and the scores of anxiety or depression symptoms, controlling for age group differences. The p-values resulting from these correlations were corrected with FDR to control for Type I errors. It should be noted that age was discretized into 5-year bins to protect participant anonymity (Babayan et al., 2019). Consequently, we report the mean age based on the minimum value of each 5-year bin for analysis.

We employed random forest regression to predict the scores of anxiety or depression symptoms, using individuals’ microstate parameters and age as input features. This approach aimed to evaluate and compare the predictive capabilities of parameters in broadband with those in specific frequency bands. A 5-fold cross-validation was conducted. In each fold, input features were standardized and converted into independent variables through principal component analysis (PCA) to reduce collinearity. Model performance was evaluated using the mean values of R-square and Mean Absolute Error (MAE). R-square, also known as the coefficient of determination, quantifies how independent variables (MS parameters and age) statistically account for the variance in dependent variable (anxiety and depression symptoms). A higher R-square indicates that the model can better explain the variability of the output with different inputs. MAE represents the average magnitude of prediction errors without direction, equally considering all discrepancies. Lower MAE signifies a model with higher predictive accuracy. We also calculated the correlation between the predictions and the true values, which served as a measure to evaluate the fitting performance of our regression prediction.

## 3 Results

### 3.1 Topographic configurations

Fig. 2 presents the microstate topographies in each condition and frequency band. With a meta-criteria method (detailed in Fig. S2), we found that the optimal number of clusters was 6 for the delta band and 4 for all other frequency bands. The results derived from the broadband EEG were consistent with the four canonical maps (Koenig et al., 2002; Michel and Koenig, 2018). However, the microstate configurations of narrowband EEG exhibited some discrepancies.

**Fig. 2.**
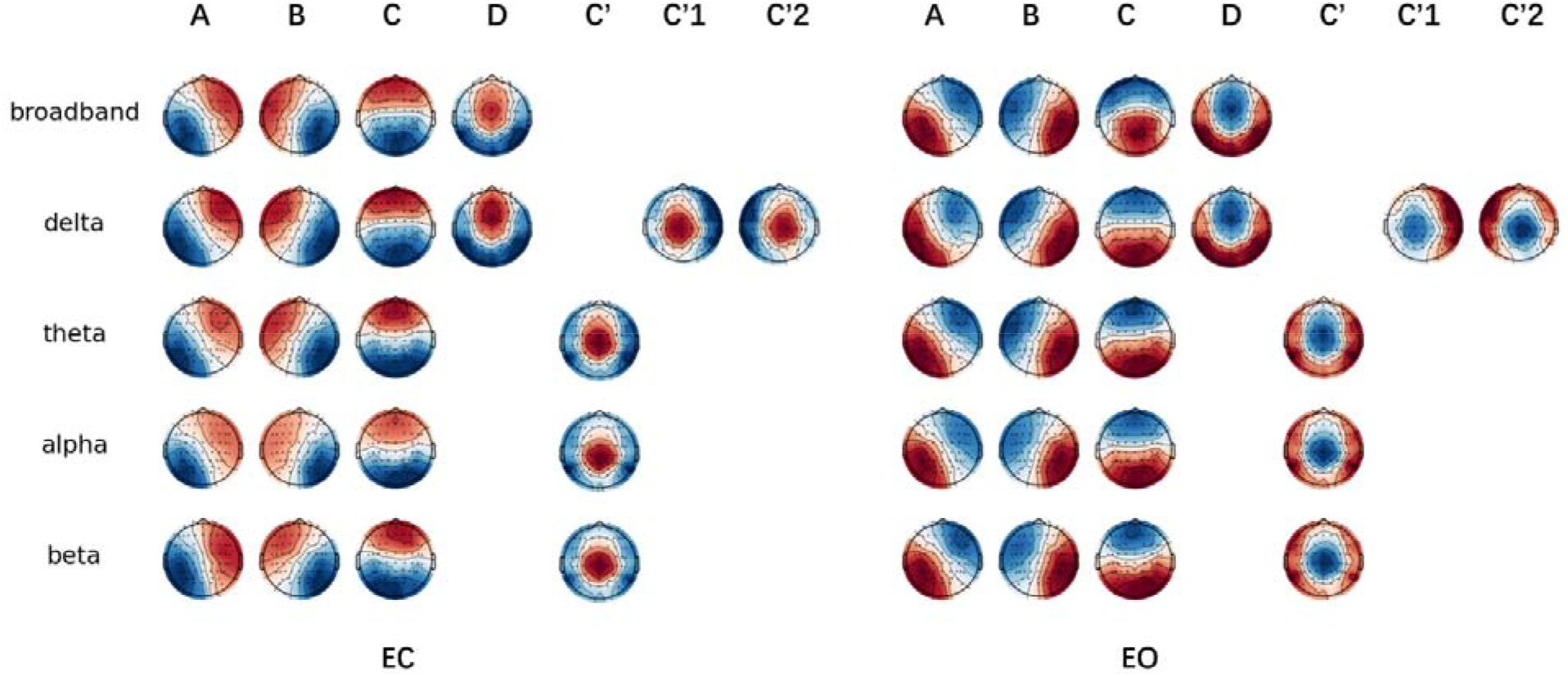
Microstate configurations across frequencies are shown for the eyes-closed (Left) and eyes-open conditions (Right). Global cluster centroids were identified from meta-criterion for each frequency band individually. Note that map polarity inversion is ignored in the classical analysis of spontaneous electroencephalography (EEG)

Within each frequency band, all 406 EEG recordings were examined. The broadband topographies together explained 62.16% (SD = 6.19%) of the GEV in the eyes-closed condition, and 57.40% (SD = 2.43%) in the eyes-open condition. In contrast, these values were higher in the delta (EC: 64.92% ± 5.88%; EO: 63.57% ± 1.58%) and alpha bands (EC: 68.45% ± 3.15%; EO: 64.10% ± 4.35%), while tended to decrease in the theta (EC: 59.36% ± 2.08%; EO: 59.39% ± 3.94%) and beta bands (EC: 51.46% ± 1.69%; EO: 53.94% ± 2.09%). This may suggest that the microstate models derived from the alpha band provided a better fit than those from the broadband.

As shown in Fig. 3, we compared the topographies between EC and EO conditions within each frequency band. All spatial correlations of the corresponding microstate were r ≥ 0.94, suggesting that topographic configurations remain stable under different conditions. We also conducted the comparison between broadband and each narrowband (as depicted in Fig. S3). Microstate topographies were designated based on canonical prototypes from previous research (Koenig et al., 2023; Zanesco, 2023), characterized by left-right orientation (A), right-left orientation (B), anterior-posterior orientation (C) and fronto-central maximum (D). Intriguingly, microstates A, B, and C exhibit consistency across frequencies (r ≥ .90). However, the cluster centroids of microstate D exhibited a posterior shift in the theta, alpha and beta bands, revealing a low correlation with that in broadband (.60 ≤ r < .87). It was thus designated as the parieto-central maximum (C’). Moreover, two additional microstates were observed within the delta band, which was defined as left parieto-central maximum (C’1) and right parieto-central maximum (C’2) due to the relatively low correlation coefficients with all broadband topographies (r ≤ .82).

**Fig. 3.**
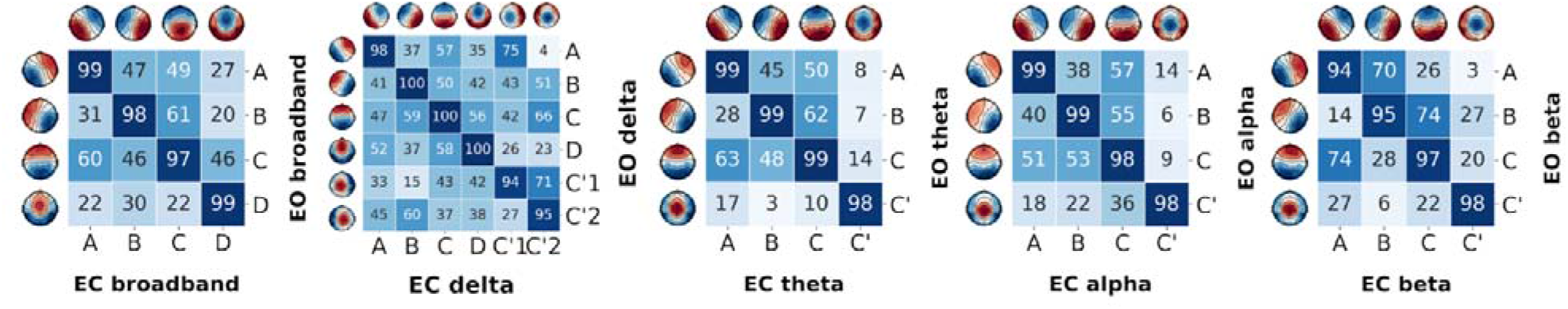
Spatial correlation of microstate topographies between eyes-closed and eyes-open conditions.

### 3.2 Frequency-specific dynamics in resting states

Paired t-tests were applied to examine the differences in microstate parameters both between and within frequencies. These analyzes involved all subjects (n = 203) and incorporated an FDR correction, as shown in Fig. 4. We also performed TESS method to calculate the neural generators from different frequency bands, which is depicted in Fig. S4-S5.

**Fig. 4.**
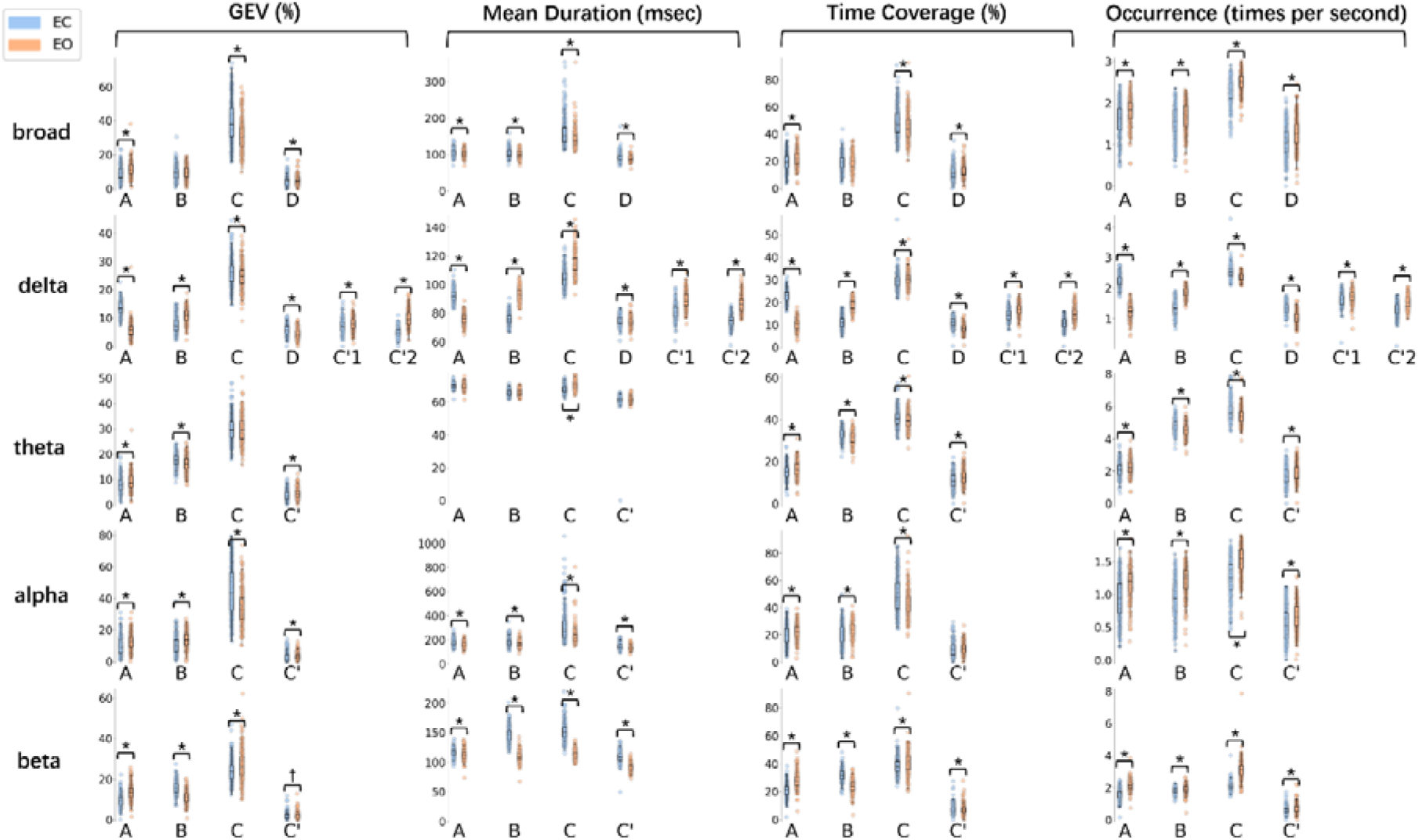
Comparison between EC and EO across all frequency bands. Significance was denoted by "*", corresponding to corrected p<0.05. Parameters with no significant difference were marked with "+".

#### Comparison between frequencies

In the delta and theta bands, there was a tendency to exhibit a shorter duration and higher occurrence compared to the broadband. Conversely, the alpha and beta bands showed an opposite trend, with a longer duration and fewer occurrence. This suggested that there were temporal differences between low (1-8 Hz) and high-frequency (8-30 Hz) components. Furthermore, specific frequency could also exhibit unique dynamics. In the delta band, many parameters were observed to be state-dependent. Compared to the broadband, microstates A and D exhibited higher GEV and occurrence under EC condition but lower under EO. Conversely, microstate B exhibited an inverse trend. In the theta band, all microstates demonstrated a higher occurrence than those in broadband, particularly in microstate B (EC: 4.81 ± 0.4, d = 8.5; EO: 4.53 ± 0.41, d = 8.31) and C (EC: 5.71 ± 0.62, d = 10.73; EO: 5.4 ± 0.48, d = 12.56). The statistical descriptions of these results can be found in Tables S1-S2.

As an examination of microstate segmentations, we conducted a comparison of the transition probabilities between broadband and each decomposed frequency band. The delta band was excluded due to variances in cluster numbers. Fig. 5 depicts the transition probability for all frequencies under both EC and EO conditions. Compared to broadband, all transition probabilities within the theta band demonstrated a significant reduction (p < .001, d range = -6.05 to -0.55), with the exception of transitions between microstates B and C. The alpha band exhibited an increase in transitions to microstates A and B (p < .008, d range = 0.47 to 1.60), and a decrease in those to C and C’ (p < .032, d range = -0.86 to - 0.22). A significant decrease in all transition probabilities was observed in the beta band (p < .001, d range = -4.86 to - 0.68). Notably, the transition probabilities to unassigned states in both theta and beta bands were significantly higher than those in broadband (p < .001, d ≥ 5.62).

**Fig. 5.**
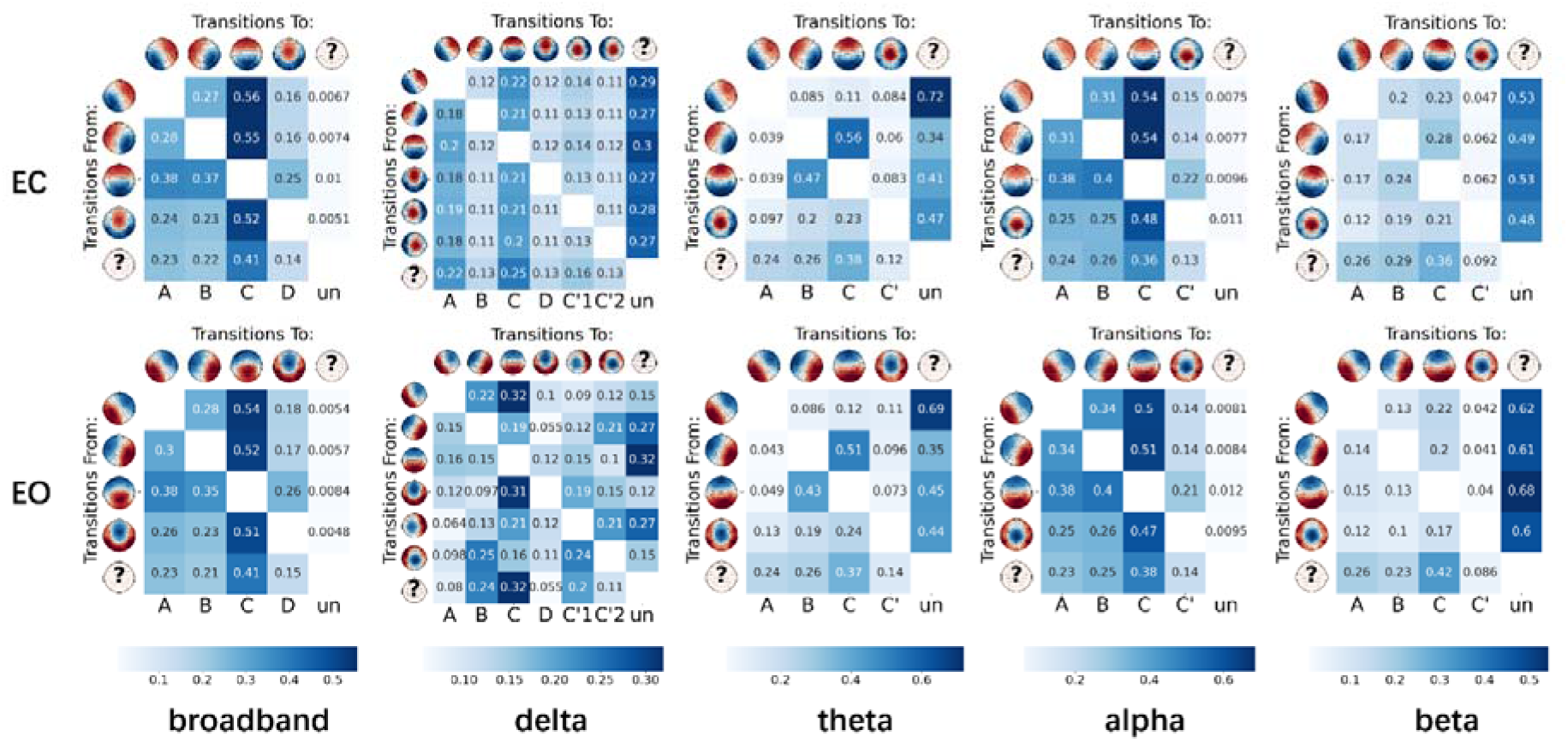
Mean Markov-chain transition probabilities for microstate configurations are shown for the eyes-closed (Top) and eyes-open conditions (Bottom). The probability of transitioning between microstates is shown for all 203 sets of microstate sequences. The ‘un’ stands for ‘unassigned’, denoting time points where the absolute spatial correlations with all topographies are below r < 0.5 threshold.

In addition, we conducted a comparison of AMI across each frequency band, as shown in Fig. S6. The mutual information was generally low, with the highest value observed between the broadband and the alpha band (EC: 9.7%; EO: 4.5%). This suggested that the microstate segmentations were comparatively independent across different frequency bands.

#### Comparison within frequencies

Resting perceptual state was an important source of change in microstate temporal dynamics. Compared to the EC condition, most microstates exhibited a shorter duration and a higher occurrence during EO. However, the low-frequency EEG bands (delta and theta) deviated from this tendency. In the delta band, microstates B (92.77 ± 4.78, d = 3.88), C (114.80 ± 7.84, d = 1.47), and C’1 (89.30 ± 5.40, d = 1.58) showed a longer duration, while the occurrence of microstates A (1.21 ± 0.23, d = -4.91) and D (1.02 ± 0.19, d = -1.55) decreased. In the theta band, duration alterations were not pronounced, while a decreased occurrence was observed in microstates B (4.53 ± 0.41, d = -0.70) and C (5.40 ± 0.48, d = -0.56).

We also investigated whether the transition probabilities differed between EC and EO conditions. The most pronounced effects were observed in the delta (d range = 0.42 to 5.88) and beta (d range = 0.33 to 3.16) bands. Moreover, when individuals opened their eyes, the transition probabilities to unassigned states tended to decrease in the delta band (p < .001, d ≤ -0.25), but increased in the beta band (p < .001, d ≥ 1.78), in comparison to when their eyes were closed.

#### Classification of EC vs. EO

In order to validate the behavioral predictive power, we applied a random forest classifier to distinguish EC vs. EO recordings across all 203 subjects, separately for each frequency bands. The feature set was divided into three parts: (i) Temporal parameters: GEV, mean duration, time coverage, and occurrence. (ii) Transition probabilities: the transitions between all microstates. (iii) All microstate parameters mentioned above.

Following a 10-fold cross-validation, it was observed that models containing features from the delta and beta bands significantly outperformed the broadband model. This was reflected in the accuracy across all feature sets: temporal parameters (delta: 99.5 ± 1.0%, beta: 98.5 ± 1.6%), transitions (delta: 100 ± 0.00%, beta: 95.6 ± 3%), and all microstate parameters (delta: 100 ± 0.0%, beta: 97.5 ± 1.9%). The AUC remained at 1.00 ± 0.00, with a slight deviation for beta band transitions (AUC = 0.98 ± 0.01). In contrast, the performance metrics derived from broadband features were considerably lower (temporal parameters: ACC = 65.3 ± 7.8%, AUC = 0.70 ± 0.08; transitions: ACC = 55.6 ± 7.9%, AUC = 0.59 ± 0.09; all microstate parameters: ACC = 67.7 ± 7.4%, AUC = 0.73 ± 0.07), with p-values of less than 0.001. These findings revealed that, narrowband microstates may provide stronger sensitivity in differentiating behavioral states in comparison to the broadband EEG. The ROC curves are depicted in Fig. 6.

**Fig. 6.**
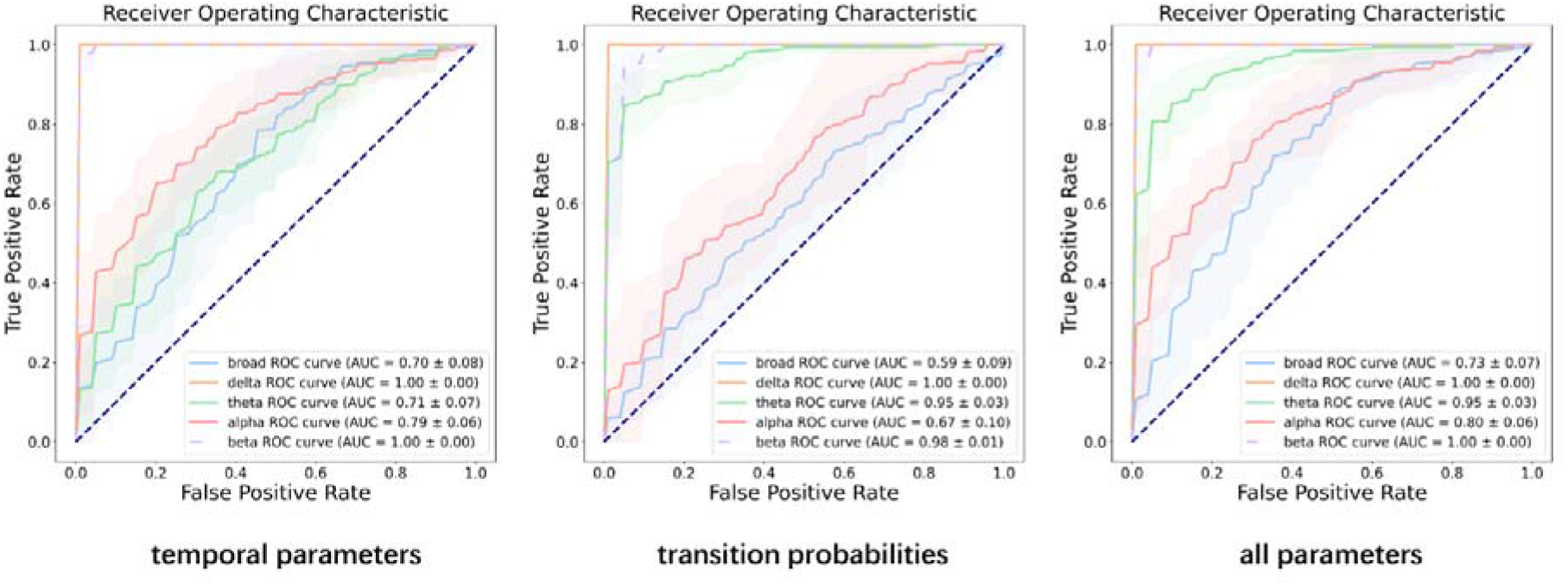
Classifying eyes-open (EO) versus eyes-closed (EC) states using microstate parameters (left: models containing four temporal parameters; middle: models containing only transition probabilities; right: models containing all microstate parameters). Binary classification performance, computed as the area under the curve (AUC) of the receiver-operating characteristic (ROC). All models shaded areas represent 95% CI. Mean AUC ± SD is reported in the legend

### 3.3 Correlation between microstate parameters and anxiety/depression symptoms

Given that age group differences were observed in both microstate parameters and predictor variables (refer to Fig. S7-S8 and Babayan et al., 2019), we controlled for age group differences in all correlations. During EC condition, five parameters within alpha band were significantly correlated with Trait-Anxiety or Hamilton Depression Scale (p ≤ .05, FDR corrected). The correlation coefficients (r) are present in Tables 1 and 2 (see Tables S3 and S4 for full results).

**Table 1.** Partial correlation analysis between temporal parameters and scales of emotional disorder (eyes-closed condition).

| temporal parameters | broadband |  | alpha |  |
| --- | --- | --- | --- | --- |
|  | Anxiety | Depression | Anxiety | Depression |
| A |  |  |  |  |
| GEV | 0.006 | 0.036 | -0.021 | 0.029 |
| Mean Duration | 0.071 | 0.007 | 0.109 | -0.002 |
| Time Coverage | -0.027 | 0.017 | -0.075 | 0.029 |
| Occurrence | -0.087 | 0.009 | -0.145 | 0.01 |
| B |  |  |  |  |
| GEV | 0.016 | 0.168 | 0.017 | 0.194* |
| Mean Duration | 0.087 | 0.134 | 0.14 | 0.133 |
| Time Coverage | 0 | 0.15 | -0.002 | 0.193* |
| Occurrence | -0.064 | 0.098 | -0.11 | 0.1 |
| C |  |  |  |  |
| GEV | 0.054 | -0.098 | 0.084 | -0.094 |
| Mean Duration | 0.118 | -0.102 | 0.179 | -0.102 |
| Time Coverage | 0.071 | -0.091 | 0.12 | -0.105 |
| Occurrence | -0.133 | 0.064 | -0.155 | 0.047 |
| D/C' |  |  |  |  |
| GEV | -0.113 | -0.007 | -0.181 | -0.049 |
| Mean Duration | -0.071 | -0.023 | -0.059 | -0.041 |
| Time Coverage | -0.119 | 0 | -0.202* | -0.034 |
| Occurrence | -0.125 | 0.009 | -0.221* | -0.025 |

Note: Significant correlations ( $p \leq .05$ , FDR control for 32 statistical tests) are denoted by an asterisk (\*).

**Table 2.**
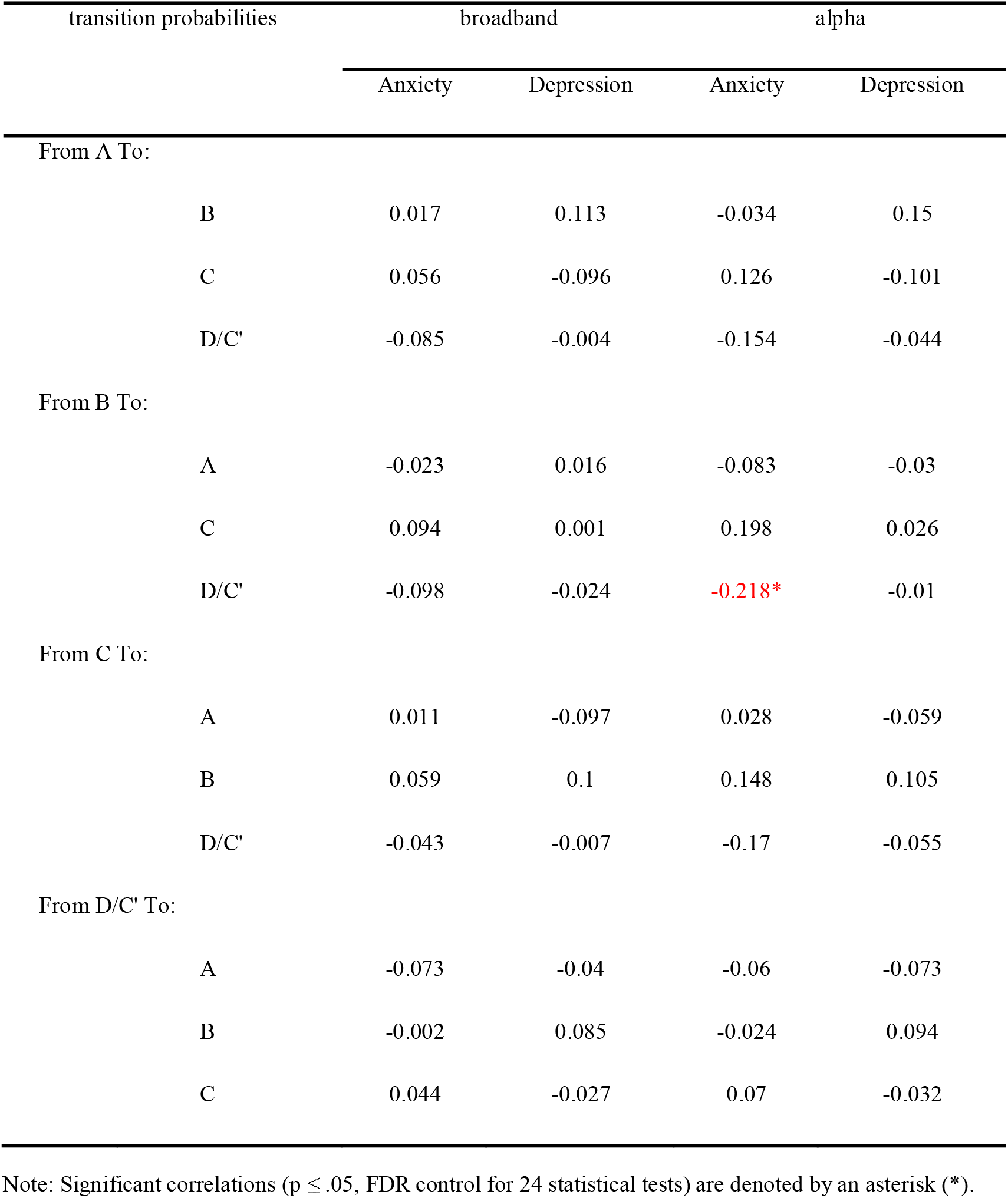
Partial correlation analysis between transition probabilities and scales of disorder (eyes-closed condition).

#### Trait-Anxiety

In the alpha band, time coverage (r = -0.202, 95% CI [-0.330, -0.070], p = 0.050) and occurrence (r = -0.221, 95% CI [-0.350, -0.090], p = 0.050) of microstate C’, as well as transition probability from B to C’ (r = -0.218, 95% CI [-0.226, -0.21], p = 0.044) exhibited a significant negative correlation with the score of trait anxiety.

#### Hamilton Depression Scale

In the alpha band, GEV (r = 0.194, 95% CI [0.060, 0.320], p = 0.050) and time coverage (r = 0.193, 95% CI [0.060, 0.320], p = 0.050) of microstate B exhibited a significant positive correlation with the score of Hamilton Scale.

These results indicate a potential for predicting an increase in the risk of anxiety and depression. Scatterplots depicting the significant correlation are shown in Fig. 7 (see Fig. S9-S12 for full results).

**Fig. 7.**
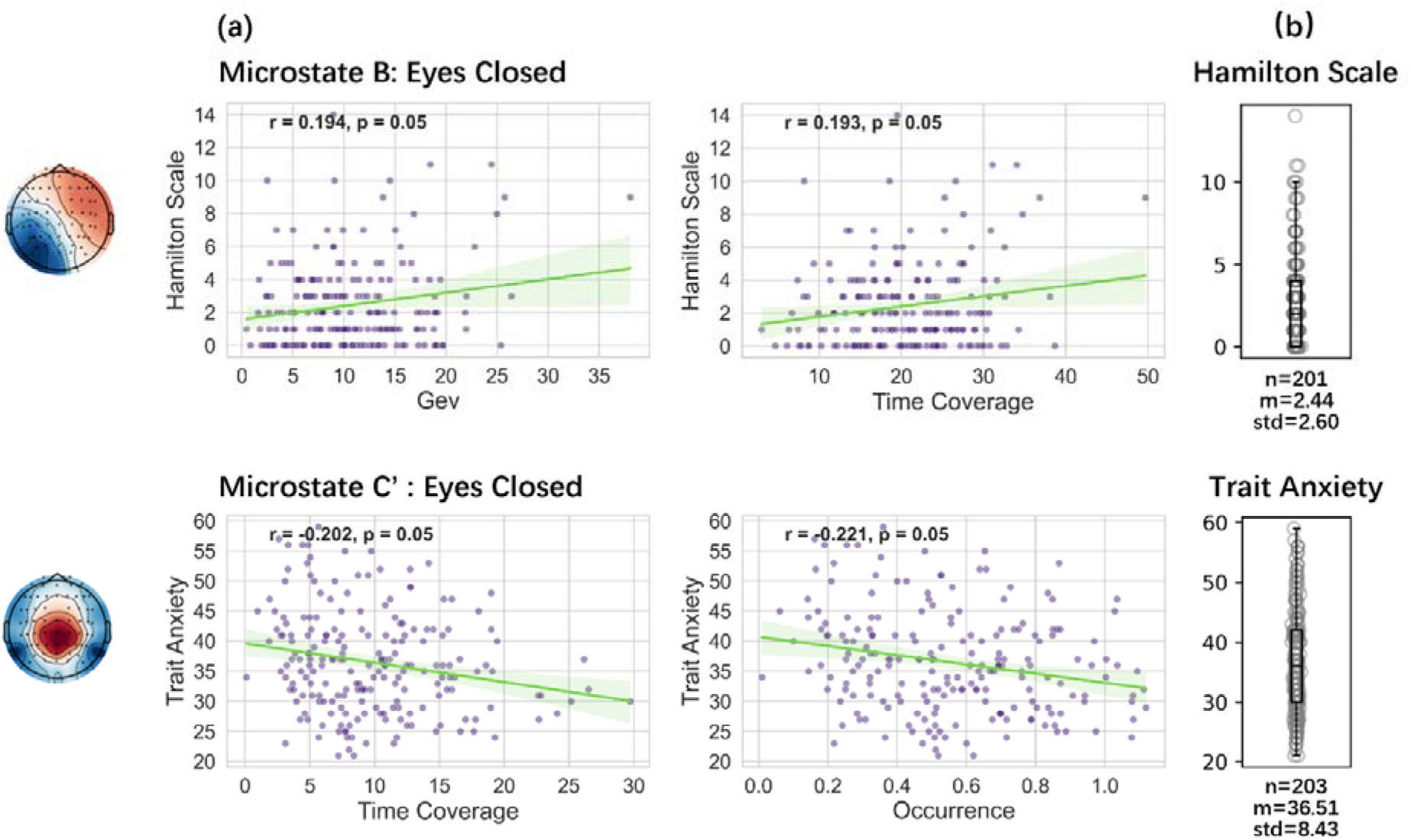
Associations between scales of emotional disorders and temporal parameters for microstates B and C’. (a) Significant correlations were found in the alpha band during eyes-closed condition: microstate B (GEV, time coverage) was positively correlated with the Hamilton Depression Scale, while microstate C’ (time coverage, occurrence) was negatively correlated with Trait Anxiety. (b) Distribution of the Hamilton Depression Scale and Trait Anxiety from the LEMON dataset

### 3.4 Predictive model for the score of anxiety/depression symptoms

In the context of predicting the severity of anxiety or depression symptoms, we hypothesized that microstate parameters in specific frequency could outperform those in broadband. The feature set was divided into three parts: (i) Temporal parameters: GEV, mean duration, time coverage, and occurrence. (ii) Transition probabilities: the transitions between all microstates. (iii) All microstate parameters mentioned above. Random forest regression was applied to predict anxiety or depression scores across all subjects, separately for EC and EO conditions. The individuals’ age was regarded as a covariate in the regression.

#### Trait-Anxiety

Following a 5-fold cross-validation, it was observed that during the EC condition, the model containing only alpha band features significantly outperformed the broadband model. This was evident in the predictive R-square values for all feature sets: temporal parameters (0.070 vs. -0.039, p < .01), transition probabilities (0.072 vs. - 0.024, p < .01) and all microstate parameters (0.100 vs. -0.026, p < .01). These results are depicted in Fig. 8 and Fig. S13 for eyes-closed condition and S14 for eyes-open condition. The eyes-closed condition yielded a much better performance than eyes-open condition.

**Fig. 8.**
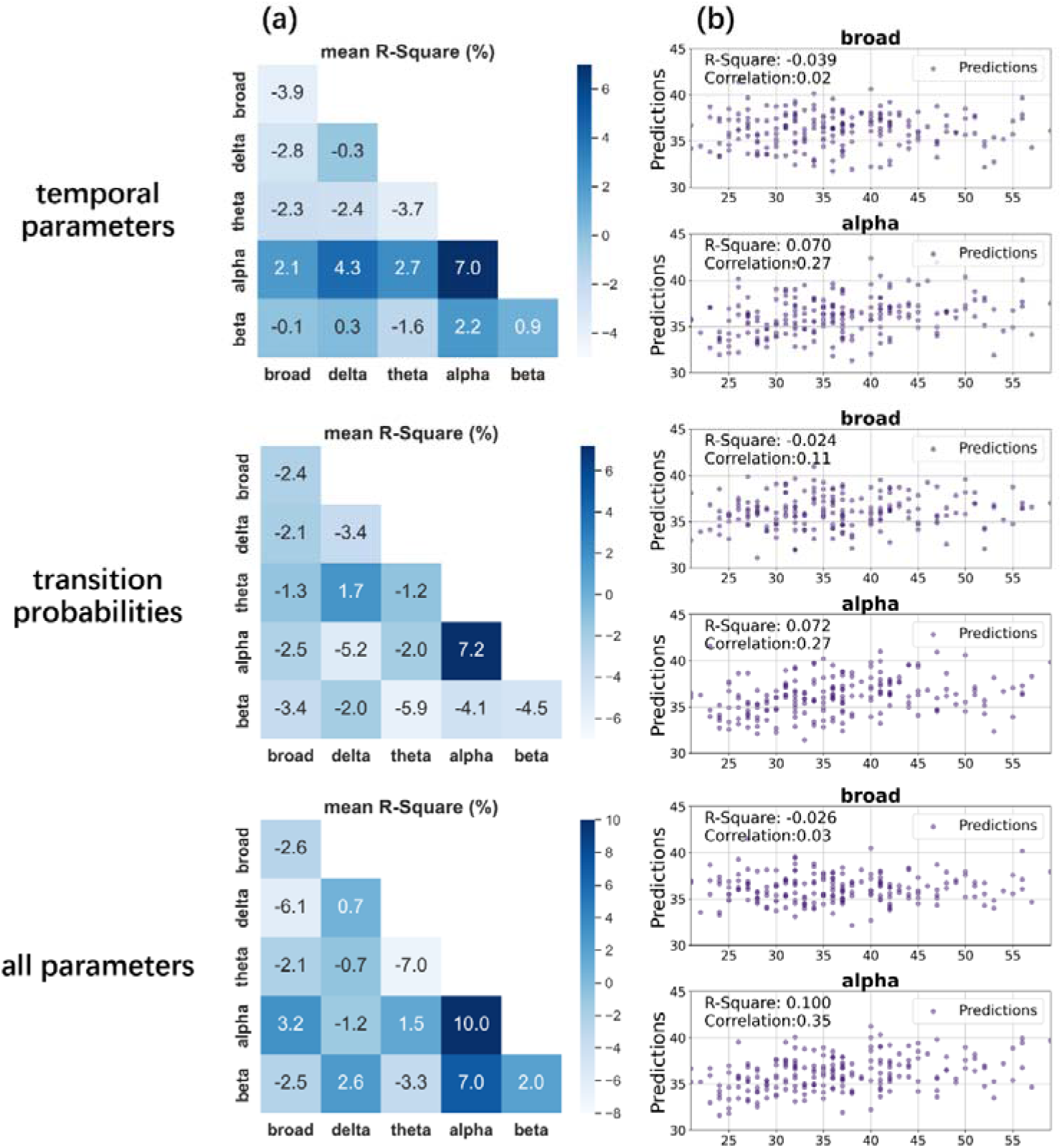
Performance of the trait anxiety prediction model (eyes-closed condition). Each grid shows the mean R-square of microstate features in one frequency band (on the diagonal) or of the combination of microstate features in two frequency bands (off the diagonal). (a) The mean R-square derived from a five-fold cross-validation procedure (top row: models containing four temporal parameters; middle row: models containing only transition probabilities; bottom row: models containing all microstate parameters). (b) Scatter plot of predictions (y-axis) versus true values (x-axis). The predictive performance significantly improves when using features derived from alpha band, as compared to using broadband features.

#### Hamilton Depression Scale

The results of depression prediction were found to be unsatisfactory, with all the R-square values falling below zero. The complete results are illustrated in Fig. S15-S16.

## 4 Discussion

The findings of this study show that, compared to broadband, narrowband microstates exhibit distinct topographic configurations and unique spatiotemporal dynamics. In the context of behavioral prediction (classifying EC vs. EO), the temporal parameters in delta and beta bands outperformed those in broadband, achieving 100% accuracy. The results of TESS method also indicated that identical topographies were derived from distinct cortical activities across different frequency bands.

Furthermore, we examined the association between narrowband microstates and anxiety or depression symptoms. In the alpha band, we discovered a positive correlation between depression and both GEV and time coverage of microstate B. Besides, time coverage and occurrence of microstate C’, as well as transitions from B to C’, were negatively correlated with anxiety. These effects were not observed in the broadband. Finally, regression analysis indicated that narrowband microstates could predict the severity of anxiety symptoms. These findings support our hypothesis that parameters in a specific frequency band might be better psychiatric predictors than those in broadband.

### 4.1 Topographic configurations of narrowband microstate

We first examined whether specific topographic microstate configurations differ across EEG frequency bands. According to the meta-criterion, the optimal number of clusters was determined as 6 for the delta band, and 4 for the other bands. In the broadband EEG, we observed four canonical maps (A-D), which aligned with previous studies (Khanna et al., 2014; Michel and Koenig, 2018). When examining spatial similarity between topographies, microstates A, B, and C were found to be consistent across different frequency bands.

These findings extend previous studies on topographies of frequency-decomposed microstates (Musaeus et al., 2020; Férat et al., 2022; Terpou et al., 2022; Xiong et al., 2023). Férat et al. (2022) and Terpou et al. (2022) fitted broadband microstate maps directly to all other frequency bands, which might neglect distinct topographic configurations in a specific frequency band. Musaeus and colleagues (2020) defined a four-cluster solution (k = 4) and reported different topographies across different EEG frequency bands. More recently, Xiong et al. (2023) defined the optimal number of clusters based on GEV and cross-validation criterion (Koenig et al., 2014). They reported that this number varies with frequency bands for task-state EEG, which enabled participants to focus on the smoking-related or paired neutral pictures. In alignment with the findings of Xiong et al., we also found different optimal cluster numbers across frequency bands in resting state EEG with a meta-criterion. This strengthens the evidence for frequency-specific cluster numbers and microstate topographies in EEG studies.

### 4.2 Frequency-specific microstate features under different perceptual states

Our findings indicated that the resting perceptual state (eyes-closed vs. eyes-open) demonstrated a frequency-specific influence on microstate features. When individuals opened their eyes, the broadband EEG microstates exhibited a shorter duration and more frequent occurrence than when their eyes were closed. These effects were consistent and more pronounced within the high-frequency EEG (alpha- and beta-band). In contrast, the low-frequency EEG (delta- and theta-band) deviated from this trend, exhibiting a less consistent variation. To further validate the applicability of narrowband microstates in the context of behavioral prediction, we employed machine learning to classify eyes-closed versus eyes-open states. The results suggest that frequency-specific microstate parameters were more accurate predictors than those derived from broadband.

These results partially conflict with those of Zanesco et al. (2020), who conducted a study of broadband EEG microstate utilizing the LEMON dataset. They reported an increased duration in microstates A, B, D and E when participants opened their eyes. In contrast, we observed a total reduction, which was in alignment with another independent work using the same dataset (Férat et al., 2022). These discrepancies may originate from the differences in EEG preprocessing and the back-fitting method. However, the decreases in GEV and duration of microstate C in the eyes-open condition were consistent across all studies, which further supports the reproducibility of microstate analysis despite the methodological differences. In addition, Férat et al. (2022) reported that when predicting eyes-closed versus eyes-open states, alpha band features were more effective than those in broadband (accuracy: 80% vs. 73%). More pronounced effects were observed in our study, with an accuracy of 100% using features from the delta or beta band. This further supports the evidence that narrowband microstates might provide stronger behavioral predictive power. The discrepancies between their study and ours may be attributed to the determination of the cluster numbers (k). Férat and colleagues fitted the five-cluster maps (k = 5) directly to all the frequency bands to obtain a common topographic configuration. In contrast, we calculated the optimal cluster number for each frequency band separately. These findings appear to support and extend the ideas of Michel and Koenig, who suggested that the optimal number of clusters should be defined in each dataset individually (Michel and Koenig, 2018). We further contend that the determination of frequency-specific cluster numbers and microstate topographies could prove to be beneficial. This approach might potentially enhance the specificity of neural markers and provide insights into brain mechanisms.

### 4.3 Transition dynamics in decomposed frequency bands

The temporal dynamics of narrowband microstates were also reflected in the transition probabilities. Among the transitions that we examined, the alpha band exhibited a similar pattern to that of the broadband. However, other frequency bands exhibited distinct transition dynamics, particularly with more frequent transitions to unassigned states. These findings extend the known properties of transition probabilities (e.g., Shen et al., 2020; Zanesco et al., 2020; Liu et al., 2023). A possible explanation could be the differential signal properties of GFP across frequency bands. Given that the periodicity of GFP is potentially determined by alpha oscillations (Koenig and Brandeis, 2016; Hari and Puce, 2017), broadband EEG microstates are inevitably driven and phase-locked by medium-high amplitude alpha rhythms (Milz et al., 2017; von Wegner et al., 2021). This might account for the observed similarity in transitions between the alpha band and broadband. Although the mutual information of the microstate sequences was relatively low, they demonstrated consistency in statistical properties. In contrast, the GFP peaks within other frequency bands provided a less definitive representation. Once the microstate topographies were fit back to continuous EEG time series, more unassigned states could be obtained, in which the voltage maps exhibit low spatial similarity (< 0.5) with all topographic configurations. This could result in a sparser microstate sequence compared to the broadband EEG. Overall, more rigorous experimental studies are needed to clarify the mechanisms underlying the representation of GFP for voltage maps in decomposed frequency bands.

### 4.4 Association between narrowband microstate and anxiety/depression symptoms

This study indicated that the abnormalities in neural activity associated with anxiety or depression were more effectively reflected in microstates derived from the alpha band, rather than those from broadband. In the alpha band, we found that microstate B was positively correlated with depression, while microstate C’ was negatively correlated with anxiety. These correlations were observed under eyes-closed condition, which ranged from 0.19 to 0.22 in magnitude. In addition, we performed random forest regression to predict the scores of anxiety or depression. The results indicated that models containing alpha band features significantly outperformed those with broadband features. These findings provided the first evidence of the narrowband microstate alterations associated with anxiety/depression.

Our findings partially align with those of (Schiller et al., 2019), who investigated the oxytocin’s effects on the microstate parameters during eyes-closed periods in 91 healthy participants. They reported that in the untreated placebo group, higher anxiety levels were associated with a shorter duration and reduced coverage of microstate D. Similarly, we found that the coverage and occurrence of microstate C’/D were negatively correlated with anxiety, an effect that was more pronounced in the alpha band than in the broadband. On the other hand, previous studies have indicated that during the resting state, all parameters of microstate B exhibit higher values in depressive patients (Gschwind et al., 2016; Yan et al., 2021; Zanesco, 2023). For instance, Yan et al., (2021) found that among the patients with depression, the duration and occurrence of microstate B significantly decreased after treatment. Once again, we observed a similar trend in the alpha band.

Furthermore, a regression analysis was performed to predict the scores of anxiety symptoms, where R-square was 0.100 for the alpha band model and -0.029 for the broadband model, respectively. Nevertheless, the predictive performance for depression was less satisfactory. A potential explanation could be the dissociated neural representations between anxiety and depression symptoms. This is in line with studies articulating circuit-based heterogeneity that relate to transdiagnostic symptoms across mood and anxiety disorders (Goldstein-Piekarski et al., 2022), and suggests that distinct types of brain circuit dysfunctions may underlie the clinical expression of depression and anxiety disorders. In addition, it is noteworthy that the EEG recordings we examined were collected from healthy adults with no history of psychiatric diseases (Babayan et al., 2019), which proved a challenge to predict emotional symptoms. This might interpret the discrepancy between our findings and those of recent studies, who suggest that the broadband EEG microstate held potential for objective diagnosis for patients with emotional disorders from healthy controls (Lei et al., 2022; Qin et al., 2022; Zhou et al., 2023).

Overall, these results confirm that frequency-decomposed (especially alpha-band) microstate analysis holds the potential to more effectively predict emotional symptoms than broadband analysis. As a result, we recommend that researchers turn their attention towards the narrowband microstate analysis. This methodology may provide a much richer description of pathological EEG dynamics, and guide new investigations into the neural substrates of psychiatric disorders.

## Acknowledgements

This work was support by National Key R&D Program of China (2023YFC2506600). We would like to thank the Mind-Body-Emotion group at the Max Planck Institute for Human Cognitive and Brain Sciences for collecting and making these data available online. We utilized the freely available Cartool software toolbox (cartoolcommunity.unige.ch) programmed by Denis Brunet, from the Functional Brain Mapping Laboratory, Geneva, Switzerland, and supported by the Center for Biomedical Imaging of Geneva and Lausanne.

## Authors contributions

Siyang Xue (Formal analysis, Software, Visualization, Writing—original draft, Writing—review & editing), Xinke Shen (Data curation, Investigation, Methodology, Writing—review & editing), Dan Zhang (Conceptualization, Validation, Writing—review & editing), Zhenhua Sang (Data curation), Qiting Long (Visualization), Sen Song (Methodology, Resources, Supervision, Writing—review & editing), and Jian Wu (Conceptualization, Funding acquisition, Investigation, Project administration)

## Data availability statement

The data that support the findings of this study are openly available in the OSF repository and can be found at: https://osf.io/u2q3r/files/osfstorage

## References

Abrol, A., Damaraju, E., Miller, R.L., Stephen, J.M., Claus, E.D., Mayer, A.R., Calhoun, V.D., 2017. Replicability of time-varying connectivity patterns in large resting state fMRI samples. Neuroimage 163, 160–176. 10.1016/j.neuroimage.2017.09.020

Al Zoubi, O., Mayeli, A., Tsuchiyagaito, A., Misaki, M., Zotev, V., Refai, H., Paulus, M., Bodurka, J., the Tulsa 1000 Investigators, Aupperle, R.L., Khalsa, S.S., Feinstein, J.S., Savitz, J., Cha, Y.-H., Kuplicki, R., Victor, T.A., 2019. EEG Microstates Temporal Dynamics Differentiate Individuals with Mood and Anxiety Disorders From Healthy Subjects. Frontiers in Human Neuroscience 13.

Anand, A., Li, Y., Wang, Y., Wu, J.W., Gao, S.J., Bukhari, L., Mathews, V.P., Kalnin, A., Lowe, M.J., 2005. Activity and connectivity of brain mood regulating circuit in depression: A functional magnetic resonance study. Biol. Psychiatry 57, 1079–1088. 10.1016/j.biopsych.2005.02.021

Babayan, A., Erbey, M., Kumral, D., Reinelt, J.D., Reiter, A.M.F., Röbbig, J., Schaare, H.L., Uhlig, M., Anwander, A., Bazin, P.-L., Horstmann, A., Lampe, L., Nikulin, V.V., Okon-Singer, H., Preusser, S., Pampel, A., Rohr, C.S., Sacher, J., Thöne-Otto, A., Trapp, S., Nierhaus, T., Altmann, D., Arelin, K., Blöchl, M., Bongartz, E., Breig, P., Cesnaite, E., Chen, S., Cozatl, R., Czerwonatis, S., Dambrauskaite, G., Dreyer, M., Enders, J., Engelhardt, M., Fischer, M.M., Forschack, N., Golchert, J., Golz, L., Guran, C.A., Hedrich, S., Hentschel, N., Hoffmann, D.I., Huntenburg, J.M., Jost, R., Kosatschek, A., Kunzendorf, S., Lammers, H., Lauckner, M.E., Mahjoory, K., Kanaan, A.S., Mendes, N., Menger, R., Morino, E., Näthe, K., Neubauer, J., Noyan, H., Oligschläger, S., Panczyszyn-Trzewik, P., Poehlchen, D., Putzke, N., Roski, S., Schaller, M.-C., Schieferbein, A., Schlaak, B., Schmidt, R., Gorgolewski, K.J., Schmidt, H.M., Schrimpf, A., Stasch, S., Voss, M., Wiedemann, A., Margulies, D.S., Gaebler, M., Villringer, A., 2019. A mind-brain-body dataset of MRI, EEG, cognition, emotion, and peripheral physiology in young and old adults. Sci Data 6, 180308. 10.1038/sdata.2018.308

Benjamini, Y., Hochberg, Y., 1995. Controlling the False Discovery Rate: A Practical and Powerful Approach to Multiple Testing. Journal of the Royal Statistical Society: Series B (Methodological) 57, 289–300. 10.1111/j.2517-6161.1995.tb02031.x

Biswal, B.B., 2012. Resting state fMRI: A personal history. NeuroImage, 20 YEARS OF fMRI 62, 938–944. 10.1016/j.neuroimage.2012.01.090

Bréchet, L., Brunet, D., Birot, G., Gruetter, R., Michel, C.M., Jorge, J., 2019. Capturing the spatiotemporal dynamics of self-generated, task-initiated thoughts with EEG and fMRI. NeuroImage 194, 82–92. 10.1016/j.neuroimage.2019.03.029

Bressler, S.L., 1995. Large-scale cortical networks and cognition. Brain Research Reviews 20, 288–304. 10.1016/0165-0173(94)00016-I

Brunet, D., Murray, M.M., Michel, C.M., 2011. Spatiotemporal Analysis of Multichannel EEG: CARTOOL. Computational Intelligence and NeuroscienceL: CIN 2011, 813870. 10.1155/2011/813870

Chivu, A., Pascal, S.A., Damborska, A., Tomescu, M.I., 2023. EEG Microstates in Mood and Anxiety Disorders: A Meta-analysis. Brain Topogr. 10.1007/s10548-023-00999-0

Custo, A., Van De Ville, D., Wells, W.M., Tomescu, M.I., Brunet, D., Michel, C.M., 2017. Electroencephalographic Resting-State Networks: Source Localization of Microstates. Brain Connect. 7, 671–682. 10.1089/brain.2016.0476

da Cruz, J.R., Favrod, O., Roinishvili, M., Chkonia, E., Brand, A., Mohr, C., Figueiredo, P., Herzog, M.H., 2020. EEG microstates are a candidate endophenotype for schizophrenia. Nature Communications 11, 3089. 10.1038/s41467-020-16914-1

Dale, A.M., Fischl, B., Sereno, M.I., 1999. Cortical Surface-Based Analysis: I. Segmentation and Surface Reconstruction. NeuroImage 9, 179–194. 10.1006/nimg.1998.0395

Diaz Hernandez, L., Rieger, K., Baenninger, A., Brandeis, D., Koenig, T., 2016. Towards Using Microstate- Neurofeedback for the Treatment of Psychotic Symptoms in Schizophrenia. A Feasibility Study in Healthy Participants. Brain Topography 29, 308–321. 10.1007/s10548-015-0460-4

Dubois, J., Galdi, P., Han, Y., Paul, L.K., Adolphs, R., 2018. Resting-State Functional Brain Connectivity Best Predicts the Personality Dimension of Openness to Experience. Personality Neuroscience 1, e6. 10.1017/pen.2018.8

Etkin, A., Wager, T.D., 2007. Functional neuroimaging of anxiety: A meta-analysis of emotional processing in PTSD, social anxiety disorder, and specific phobia. Am. J. Psychiat. 164, 1476–1488. 10.1176/appi.ajp.2007.07030504

Férat, V., Seeber, M., Michel, C.M., Ros, T., 2022. Beyond broadband: Towards a spectral decomposition of electroencephalography microstates. Human Brain Mapping 43, 3047–3061. 10.1002/hbm.25834

Feurer, C., Jimmy, J., Chang, F., Langenecker, S.A., Phan, K.L., Ajilore, O., Klumpp, H., 2021. Resting state functional connectivity correlates of rumination and worry in internalizing psychopathologies. Depress. Anxiety 38, 488–497. 10.1002/da.23142

Gibbons, R.D., Hedeker, D.R., Davis, J.M., 1993. Estimation of Effect Size from a Series of Experiments Involving Paired Comparisons. Journal of Educational Statistics 18, 271–279. 10.2307/1165136

Goldstein-Piekarski, A.N., Ball, T.M., Samara, Z., Staveland, B.R., Keller, A.S., Fleming, S.L., Grisanzio, K.A., Holt-Gosselin, B., Stetz, P., Ma, J., Williams, L.M., 2022. Mapping Neural Circuit Biotypes to Symptoms and Behavioral Dimensions of Depression and Anxiety. Biological Psychiatry 91, 561–571. 10.1016/j.biopsych.2021.06.024

Gramfort, A., Luessi, M., Larson, E., Engemann, D.A., Strohmeier, D., Brodbeck, C., Goj, R., Jas, M., Brooks, T., Parkkonen, L., Hämäläinen, M., 2013. MEG and EEG data analysis with MNE-Python. Front. Neurosci. 7, 70133. 10.3389/fnins.2013.00267

Grieder, M., Koenig, T., Kinoshita, T., Utsunomiya, K., Wahlund, L.-O., Dierks, T., Nishida, K., 2016. Discovering EEG resting state alterations of semantic dementia. Clinical Neurophysiology 127, 2175–2181. 10.1016/j.clinph.2016.01.025

Gschwind, M., Hardmeier, M., Van De Ville, D., Tomescu, M.I., Penner, I.-K., Naegelin, Y., Fuhr, P., Michel, C.M., Seeck, M., 2016. Fluctuations of spontaneous EEG topographies predict disease state in relapsing-remitting multiple sclerosis. Neuroimage Clin 12, 466–477. 10.1016/j.nicl.2016.08.008

Hari, R., Puce, A., 2017. MEG-EEG Primer. MEG-EEG Primer.

Khanna, A., Pascual-Leone, A., Farzan, F., 2014. Reliability of Resting-State Microstate Features in Electroencephalography. PLoS ONE 9, e114163. 10.1371/journal.pone.0114163

Khanna, A., Pascual-Leone, A., Michel, C.M., Farzan, F., 2015. Microstates in resting-state EEG: Current status and future directions. Neuroscience & Biobehavioral Reviews 49, 105–113. 10.1016/j.neubiorev.2014.12.010

Kim, K., Duc, N.T., Choi, M., Lee, B., 2021. EEG microstate features for schizophrenia classification. PLOS ONE 16, e0251842. 10.1371/journal.pone.0251842

Koenig, T., Brandeis, D., 2016. Inappropriate assumptions about EEG state changes and their impact on the quantification of EEG state dynamics. NeuroImage 125, 1104–1106. 10.1016/j.neuroimage.2015.06.035

Koenig, T., Diezig, S., Kalburgi, S.N., Antonova, E., Artoni, F., Brechet, L., Britz, J., Croce, P., Custo, A., Damborská, A., Deolindo, C., Heinrichs, M., Kleinert, T., Liang, Z., Murphy, M.M., Nash, K., Nehaniv, C., Schiller, B., Smailovic, U., Tarailis, P., Tomescu, M., Toplutaş, E., Vellante, F., Zanesco, A., Zappasodi, F., Zou, Q., Michel, C.M., 2023. EEG-Meta-Microstates: Towards a More Objective Use of Resting-State EEG Microstate Findings Across Studies. Brain Topogr. 10.1007/s10548-023-00993-6

Koenig, T., Prichep, L., Lehmann, D., Sosa, P.V., Braeker, E., Kleinlogel, H., Isenhart, R., John, E.R., 2002. Millisecond by Millisecond, Year by Year: Normative EEG Microstates and Developmental Stages. NeuroImage 16, 41–48. 10.1006/nimg.2002.1070

Koenig, T., Stein, M., Grieder, M., Kottlow, M., 2014. A Tutorial on Data-Driven Methods for Statistically Assessing ERP Topographies. Brain Topogr 27, 72–83. 10.1007/s10548-013-0310-1

Krug, S.E., 1976. Handbook for the IPAT anxiety scale. Institute for Personality and Ability Testing, Champaign, Ill.

Lassi, M., Fabbiani, C., Mazzeo, S., Burali, R., Vergani, A.A., Giacomucci, G., Moschini, V., Morinelli, C., Emiliani, F., Scarpino, M., Bagnoli, S., Ingannato, A., Nacmias, B., Padiglioni, S., Micera, S., Sorbi, S., Grippo, A., Bessi, V., Mazzoni, A., 2023. Degradation of EEG microstates patterns in subjective cognitive decline and mild cognitive impairment: Early biomarkers along the Alzheimer’s Disease continuum? NeuroImage: Clinical 38, 103407. 10.1016/j.nicl.2023.103407

Lehmann, D., 1971. Multichannel topography of human alpha EEG fields. Electroencephalography and Clinical Neurophysiology 31, 439–449. 10.1016/0013-4694(71)90165-9

Lehmann, D., Ozaki, H., Pal, I., 1987. EEG alpha map series: brain micro-states by space-oriented adaptive segmentation. Electroencephalography and Clinical Neurophysiology 67, 271–288. 10.1016/0013-4694(87)90025-3

Lei, L., Liu, Z., Zhang, Y., Guo, M., Liu, P., Hu, X., Yang, C., Zhang, A., Sun, N., Wang, Y., Zhang, K., 2022. EEG microstates as markers of major depressive disorder and predictors of response to SSRIs therapy. Progress in Neuro-Psychopharmacology and Biological Psychiatry 116, 110514. 10.1016/j.pnpbp.2022.110514

Li, J., Li, N., Shao, X., Chen, J., Hao, Y., Li, X., Hu, B., 2022. Altered Brain Dynamics and Their Ability for Major Depression Detection Using EEG Microstates Analysis. IEEE Trans. Affect. Comput. 14, 2116–2126. 10.1109/TAFFC.2021.3139104

Liu, J., Hu, X., Shen, X., Lv, Z., Song, S., Zhang, D., 2023. The EEG microstate representation of discrete emotions. International Journal of Psychophysiology 186, 33–41. 10.1016/j.ijpsycho.2023.02.002

Martınez-Montes, E., Valdés-Sosa, P.A., Miwakeichi, F., Goldman, R.I., Cohen, M.S., 2004. Concurrent EEG/fMRI analysis by multiway Partial Least Squares. NeuroImage 22, 1023–1034. 10.1016/j.neuroimage.2004.03.038

Michel, C.M., Koenig, T., 2018. EEG microstates as a tool for studying the temporal dynamics of whole-brain neuronal networks: A review. NeuroImage, Brain Connectivity Dynamics 180, 577–593. 10.1016/j.neuroimage.2017.11.062

Michel, C.M., Murray, M.M., Lantz, G., Gonzalez, S., Spinelli, L., Grave de Peralta, R., 2004. EEG source imaging. Clinical Neurophysiology 115, 2195–2222. 10.1016/j.clinph.2004.06.001

Milz, P., Pascual-Marqui, R.D., Achermann, P., Kochi, K., Faber, P.L., 2017. The EEG microstate topography is predominantly determined by intracortical sources in the alpha band. Neuroimage 162, 353–361. 10.1016/j.neuroimage.2017.08.058

Murphy, M., Whitton, A.E., Deccy, S., Ironside, M.L., Rutherford, A., Beltzer, M., Sacchet, M., Pizzagalli, D.A., 2020. Abnormalities in electroencephalographic microstates are state and trait markers of major depressive disorder. Neuropsychopharmacol. 45, 2030–2037. 10.1038/s41386-020-0749-1

Murray, M.M., Brunet, D., Michel, C.M., 2008. Topographic ERP Analyses: A Step-by-Step Tutorial Review. Brain Topography 20, 249–64. 10.1007/s10548-008-0054-5

Musaeus, C.S., Engedal, K., Hogh, P., Jelic, V., Khanna, A.R., Kjaer, T.W., Morup, M., Naik, M., Oeksengaard, A.-R., Santarnecchi, E., Snaedal, J., Wahlund, L.-O., Waldemar, G., Andersen, B.B., 2020. Changes in the left temporal microstate are a sign of cognitive decline in patients with Alzheimer’s disease. Brain Behav. 10, e01630. 10.1002/brb3.1630

Pascual-Marqui, R.D., Michel, C.M., Lehmann, D., 1995. Segmentation of brain electrical activity into microstates: model estimation and validation. IEEE Transactions on Biomedical Engineering 42, 658–665. 10.1109/10.391164

Pizzagalli, D.A., 2011. Frontocingulate Dysfunction in Depression: Toward Biomarkers of Treatment Response. Neuropsychopharmacology 36, 183–206. 10.1038/npp.2010.166

Preti, M.G., Bolton, T.A.W., Van De Ville, D., 2017. The dynamic functional connectome: State-of-the-art and perspectives. Neuroimage 160, 41–54. 10.1016/j.neuroimage.2016.12.061

Qin, X., Xiong, J., Cui, R., Zou, G., Long, C., Lei, X., 2022. EEG microstate temporal Dynamics Predict depressive symptoms in College Students. Brain Topogr 35, 481–494. 10.1007/s10548-022-00905-0

Raichle, M.E., 2015. The Brain’s Default Mode Network, in: Hyman, S.E. (Ed.), Annual Review of Neuroscience, VOL 38. Annual Reviews, Palo Alto, pp. 433–447. 10.1146/annurev-neuro-071013-014030

Raichle, M.E., 2010. Two views of brain function. Trends in Cognitive Sciences 14, 180–190. 10.1016/j.tics.2010.01.008

Schiller, B., Koenig, T., Heinrichs, M., 2019. Oxytocin modulates the temporal dynamics of resting EEG networks. Sci Rep 9, 13418. 10.1038/s41598-019-49636-6

Shen, X., Hu, X., Liu, S., Song, S., Zhang, D., 2020. Exploring EEG microstates for affective computing: decoding valence and arousal experiences during video watching, in: 42nd Annual International Conferences Of The Ieee Engineering In Medicine And Biology Society: Enabling Innovative Technologies For Global Healthcare EMBC’20, IEEE Engineering in Medicine and Biology Society Conference Proceedings. Presented at the 42nd Annual International Conference of the IEEE-Engineering-in-Medicine-and-Biology-Society (EMBC), IEEE, New York, pp. 841–846.

Spielberger, C., Gorsuch, R., Lushene, R., 1970. Manual for the State-Trait Anxiety Inventory.

Spielberger, C.D., Reheiser, E.C., 2009. Assessment of Emotions: Anxiety, Anger, Depression, and Curiosity. Applied Psychology: Health and Well-Being 1, 271–302. 10.1111/j.1758-0854.2009.01017.x

Sylvester, C.M., Corbetta, M., Raichle, M.E., Rodebaugh, T.L., Schlaggar, B.L., Sheline, Y.I., Zorumski, C.F., Lenz, E.J., 2012. Functional network dysfunction in anxiety and anxiety disorders. Trends Neurosci. 35, 527–535. 10.1016/j.tins.2012.04.012

Tadel, F., Baillet, S., Mosher, J.C., Pantazis, D., Leahy, R.M., 2011. Brainstorm: A User-Friendly Application for MEG/EEG Analysis. Computational Intelligence and Neuroscience 2011, e879716. 10.1155/2011/879716

Taylor, J.A., 1953. A personality scale of manifest anxiety. The Journal of Abnormal and Social Psychology 48, 285–290. 10.1037/h0056264

Terpou, B.A., Shaw, S.B., Theberge, J., Ferat, V., Michel, C.M., McKinnon, M.C., Lanius, R.A., Ros, T., 2022. Spectral decomposition of EEG microstates in post-traumatic stress disorder. NeuroImage-Clin. 35, 103135. 10.1016/j.nicl.2022.103135

Thirioux, B., Langbour, N., Bokam, P., Renaudin, L., Wassouf, I., Harika-Germaneau, G., Jaafari, N., 2023. Microstates imbalance is associated with a functional dysregulation of the resting-state networks in obsessive–compulsive disorder: a high-density electrical neuroimaging study using the TESS method. Cerebral Cortex 33, 2593–2611. 10.1093/cercor/bhac229

Tomescu, M.I., Rihs, T.A., Becker, R., Britz, J., Custo, A., Grouiller, F., Schneider, M., Debbané, M., Eliez, S., Michel, C.M., 2014. Deviant dynamics of EEG resting state pattern in 22q11.2 deletion syndrome adolescents: A vulnerability marker of schizophrenia? Schizophrenia Research 157, 175–181. 10.1016/j.schres.2014.05.036

Trevor Hastie, Robert Tibshirani, Jerome Friedman, 2009. The Elements of Statistical Learning.

Van De Ville, D., Britz, J., Michel, C.M., Logothetis, N.K., 2010. EEG microstate sequences in healthy humans at rest reveal scale-free dynamics. Proc Natl Acad Sci U S A 107, 18179–18184. 10.1073/pnas.1007841107

Vinh, N.X., Epps, J., Bailey, J., 2010. Information Theoretic Measures for Clusterings Comparison: Variants, Properties, Normalization and Correction for Chance.

von Wegner, F., Bauer, S., Rosenow, F., Triesch, J., Laufs, H., 2021. EEG microstate periodicity explained by rotating phase patterns of resting-state alpha oscillations. NeuroImage 224, 117372. 10.1016/j.neuroimage.2020.117372

Wang, S., Zhao, Y., Zhang, L., Wang, Xu, Wang, Xiuli, Cheng, B., Luo, K., Gong, Q., 2019. Stress and the brain: Perceived stress mediates the impact of the superior frontal gyrus spontaneous activity on depressive symptoms in late adolescence. Hum Brain Mapp 40, 4982–4993. 10.1002/hbm.24752

Xiong, X., Feng, J., Zhang, Y., Wu, D., Yi, S., Wang, C., Liu, R., He, J., 2023. Improved HHT-microstate analysis of EEG in nicotine addicts. Front. Neurosci. 17, 1174399. 10.3389/fnins.2023.1174399

Xu, J., Van Dam, N.T., Feng, C., Luo, Y., Ai, H., Gu, R., Xu, P., 2019. Anxious brain networks: A coordinate-based activation likelihood estimation meta-analysis of resting-state functional connectivity studies in anxiety. Neurosci. Biobehav. Rev. 96, 21–30. 10.1016/j.neubiorev.2018.11.005

Yan, D., Liu, J., Liao, M., Liu, B., Wu, S., Li, X., Li, H., Ou, W., Zhang, L., Li, Z., Zhang, Y., Li, L., 2021. Prediction of Clinical Outcomes With EEG Microstate in Patients With Major Depressive Disorder. Front. Psychiatry 12, 695272. 10.3389/fpsyt.2021.695272

Zanesco, A.P., 2023. Normative Temporal Dynamics of Resting EEG Microstates. Brain Topography. 10.1007/s10548-023-01004-4

Zhou, D.-D., Peng, X.-Y., Zhao, L., Ma, L.-L., Hu, J.-H., Jiang, Z.-H., He, X.-Q., Wang, W., Chen, R., Kuang, L., 2023. Neurophysiological biomarkers for depression classification: Utilizing microstate k-mers and a bag-of-words model. J. Psychiatr. Res. 165, 197–204. 10.1016/j.jpsychires.2023.07.021

