## Supplementary material for "Unveiling Frequency-Specific Microstate Correlates of Anxiety and Depression Symptoms"

**1 Source localization**

We utilized TESS method (Custo et al., 2014, 2017) to estimate the neural generators of microstate (MS) topographies. TESS relies upon two general linear models (GLM). The first GLM is used to identify significant states and their temporal regressors of microstates, which is fitted to each participant’s continuous EEG. The second GLM computes corresponding sources from these temporal regressors.

Source localization was achieved through dSPM (dynamic statistical parametric mapping), a noise-normalized linear estimates method (Dale et al., 1999). We applied dSPM to each time point of the individual preprocessed EEG data and estimated the time course of the current density of the 17001 solution points (Michel et al., 2004) via the Brainstorm (Tadel et al., 2011).

This time course was fitted with the temporal regressors through GLM, allowing us to estimate the coefficient for each solution point and each microstate. Finally, we performed bootstrapping (Trevor Hastie et al., 2009) to examine significantly active voxels for each estimated activity map, thus determining EEG based resting-state networks (eRSNs). The significant estimated betas give us the location and amplitude of the generators of each microstate, as shown in Fig. S1. More details can be found in (Custo et al., 2017).

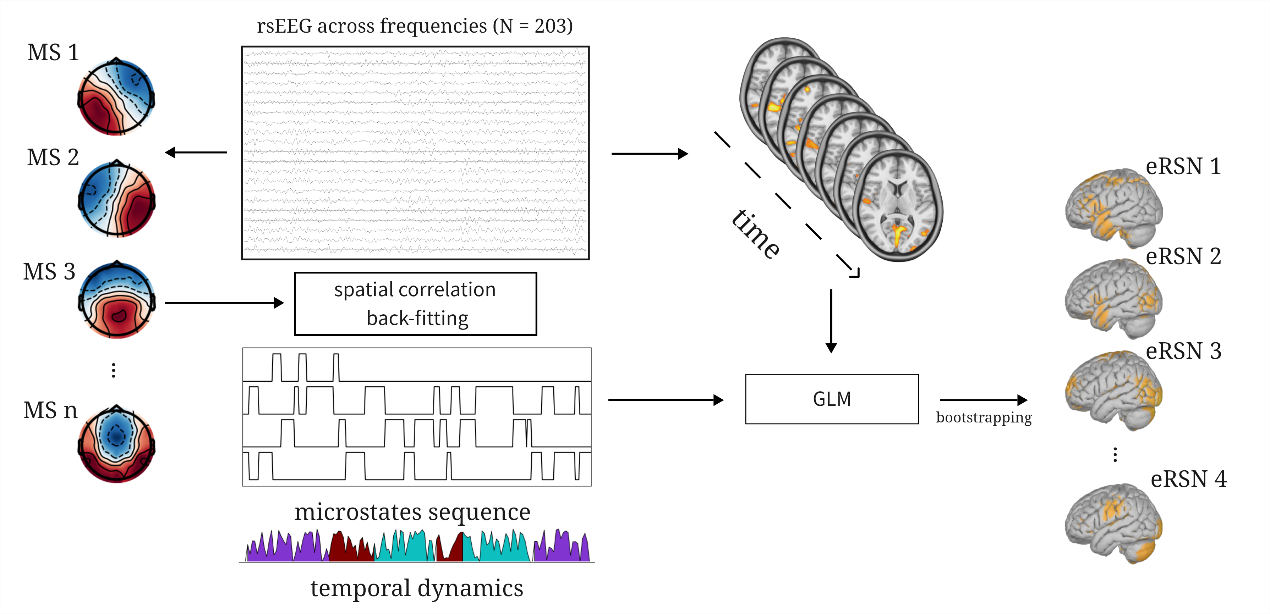

Fig. S1 Methodological schema of TESS method.

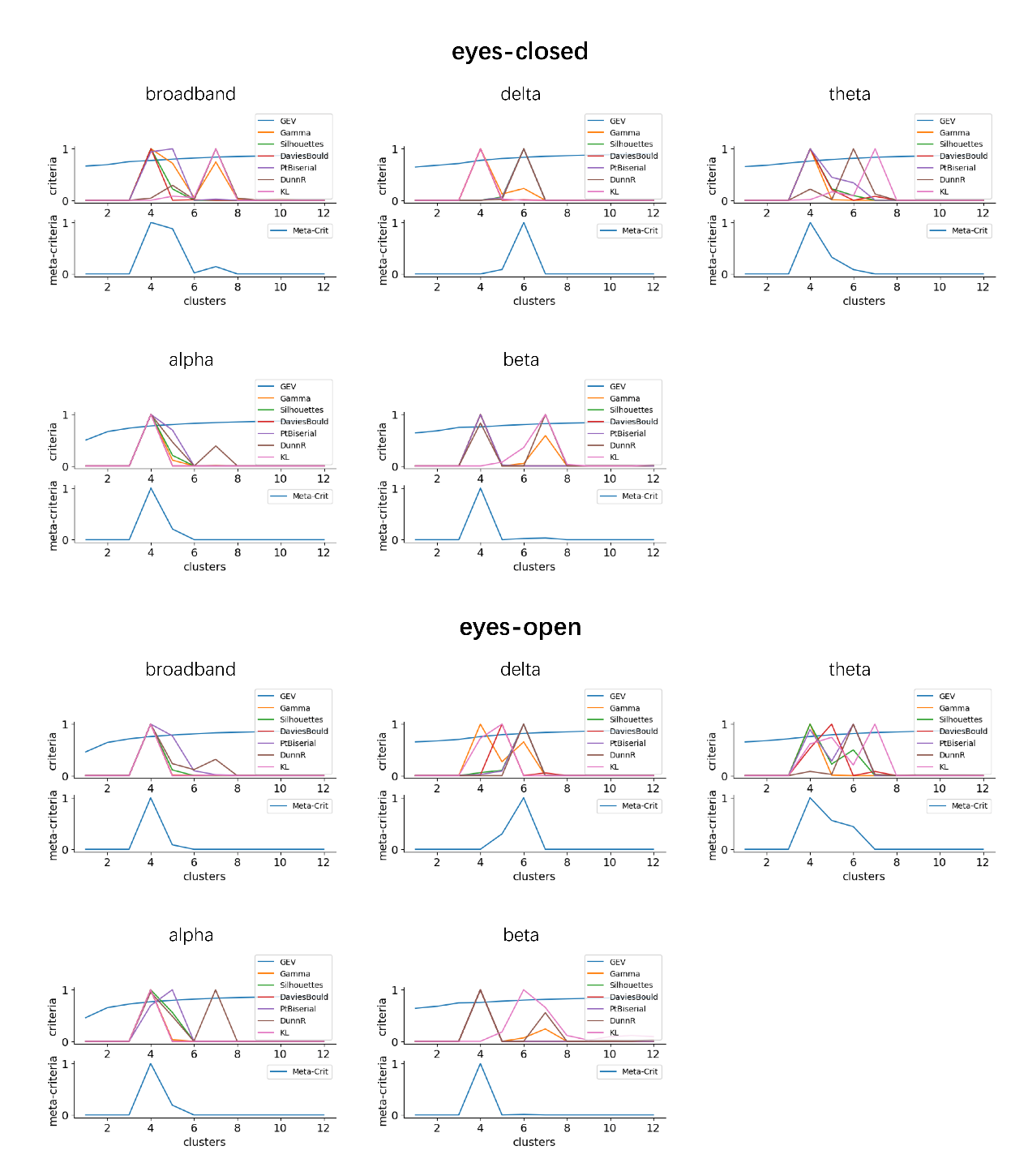

Fig. S2 The results of meta-criterion for all frequencies and conditions (7 criteria). The optimal number of clusters is defined with the peak of the meta-criterion.

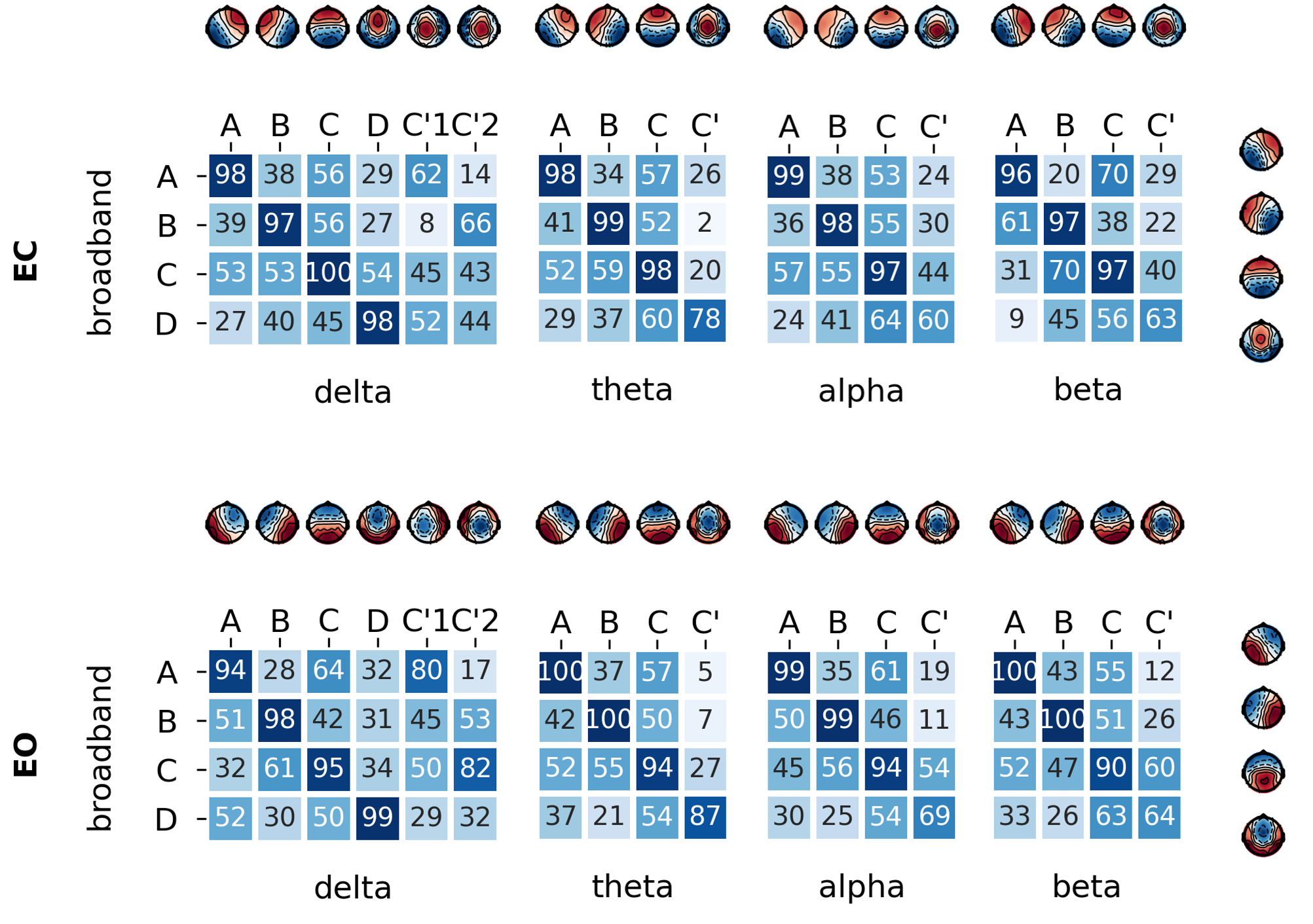

Fig. S3 Spatial correlations are depicted between each frequency and broadband for eyes-closed (EC) and eyes-open (EO) condition. Polarity is ignored and only the spatial configuration is considered.

**2 Frequency-specific neural generators**

In this study, we employed the z-score, a statistical metric obtained through bootstrapping, to offer a robust estimation of neural activity within the population. Results revealed that neural activity was more **intense** in the broadband compared to the narrowband. Therefore, we defined the thresholds to denote an active area as z > 3 for broadband and z > 2 for narrowband. Results are provided in Fig. S5 and Fig. S6.

In broadband, eRSN for microstate A was detected in the parietal and occipital lobes (BA7, 17-19). Microstate B activated in the cuneus and precuneus (BA7, 17-19). Microstate C was primarily located in the right temporal lobe (BA20-22, 39, 41, 42), precentral and postcentral gyrus (BA1-4). Microstate D was mainly active in the left temporal lobe (BA21, 38, 43) and the posterior cingulate cortex (BA31). However, the source activity in narrowband exhibited significant variation compared to broadband. For instance, in the delta band, eRSN for microstate A was concentrated in the left middle temporal gyrus, left superior temporal gyrus, and left superior frontal gyrus (BA8, 21, 22). In the theta band, it appeared in the left superior occipital gyrus, right superior parietal gyrus, middle occipital gyrus, and precuneus (BA5, 7, 18, 19). In the alpha band, it was observed in the cuneus, precuneus, lingual gyrus, superior occipital gyrus, and superior parietal gyrus (BA5, 7, 17-19). In the beta band, it was concentrated in the left temporal lobe, left orbitofrontal cortex, pars opercularis, pars triangularis, middle frontal gyrus, and subcentral area (BA11, 20, 43-46). Furthermore, among all the topographies, D (C’) of the low-frequency bands (delta, theta) showed a reduced level of activation, which might be associated with its lower global explained variance (GEV).

Consistent with past research, our findings indicate that activity within the alpha band was primarily located in the parietal and occipital lobes, while the delta band demonstrated a concentration in the frontal lobe (Goldman et al., 2001; Groppe et al., 2013; Mantini et al., 2007; Mellem et al., 2017). These results further reveal that identical microstates originated from different generators across various frequencies. Therefore, the results lend support to the idea that same EEG topographies do not necessarily imply identical neural generators (Michel et al., 2004). Microstate in different frequency band might reflect different neural processes in the brain.

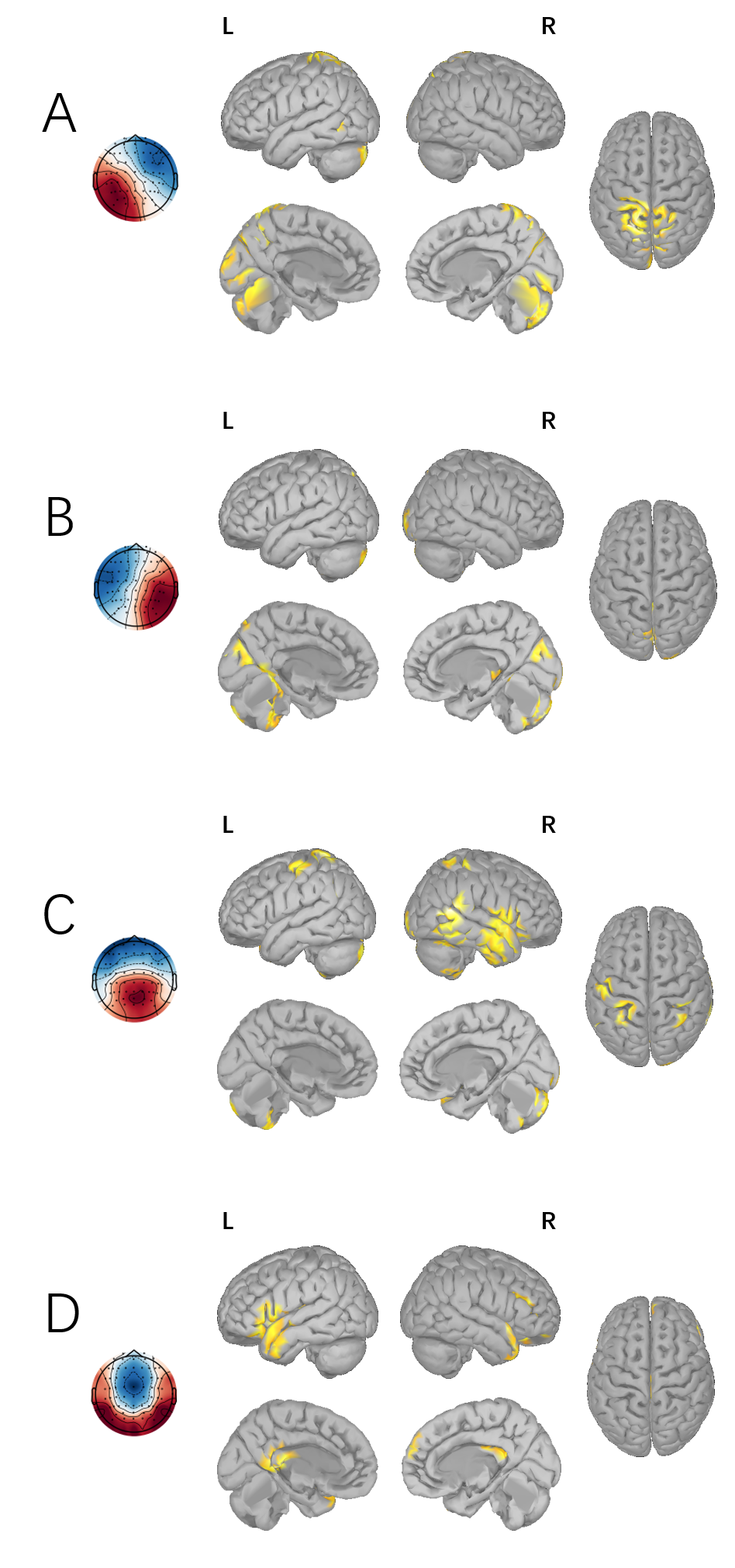

Fig. S4 The estimated RSNs (A–D) of broadband. The z scores resulting from bootstrapping (p < .005) are thresholded at z > 3.

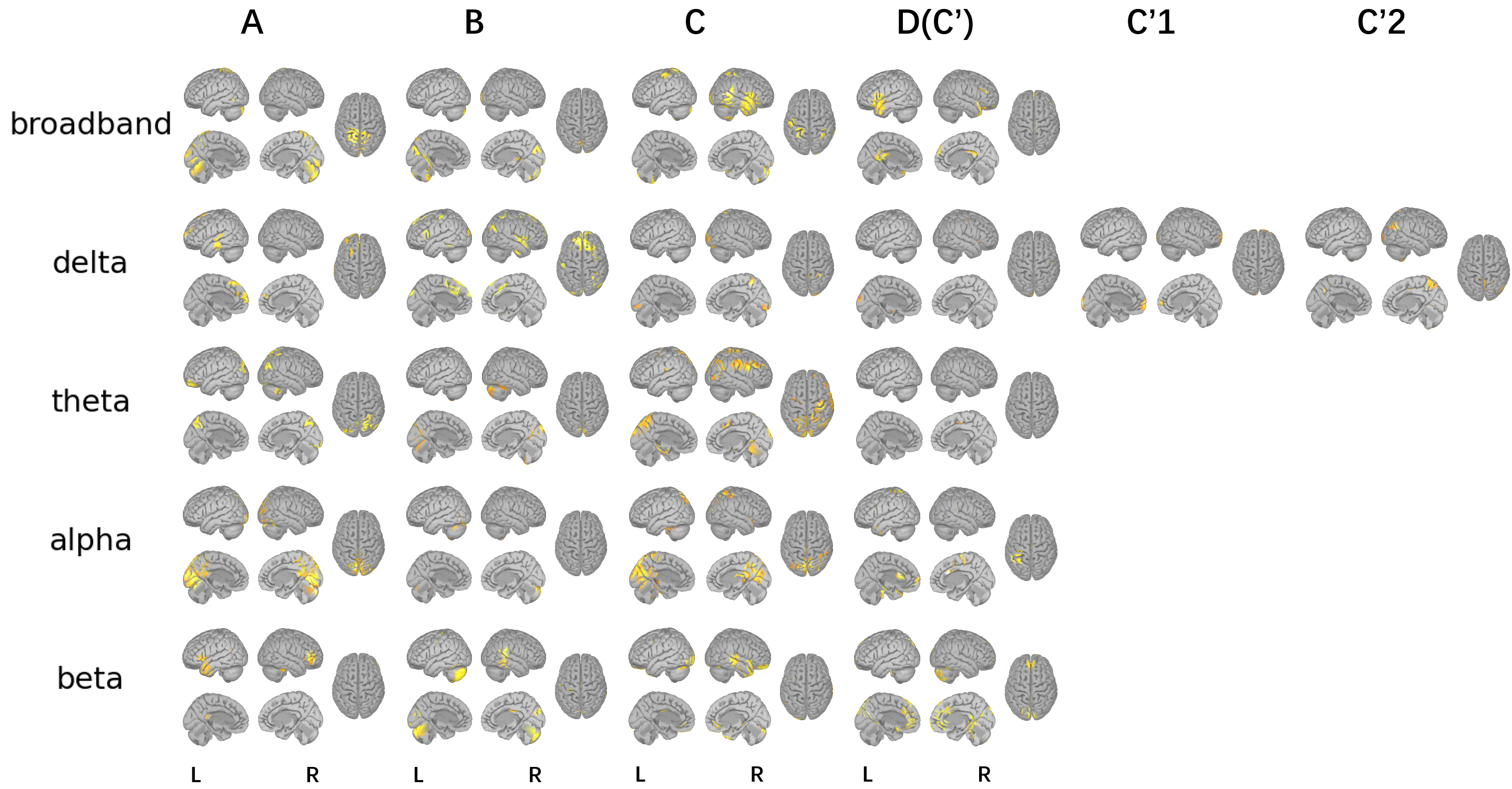

Fig. S5 The estimated RSNs for each frequency (eyes-closed condition). The z scores are thresholded at z > 3 for broadband and z > 2 for narrowband.

**3 AMI Between Different Frequency Bands**

Fig. S4 shows that during EC condition, the AMI between broadband and narrowband segmentation for delta, theta, alpha, and beta were 1.1%, 1.0%, 9.7%, and 0.5% respectively. It is evident that the mutual information of the decomposed frequency bands with broadband is quite low, which suggest independence between broadband and narrowband microstate segmentations.

Within the narrowband, there is a certain correlation between delta and theta, the two low-frequency bands (EC: s = 2.3%, EO: s = 2.4%), but their correlation with high frequency (alpha, beta) is very low (s $\leq$ 0.3%). When the experimental condition transitions from EC to EO, the information shared with broadband increases for the delta band (from s = 1.1% to s = 2.7%), but decreases for the alpha band (from s = 9.7% to s = 4.5%). The latter aligns with expectations as it is well known that when the eyes are closed, alpha oscillations significantly increase, amplifying their contribution to the broadband signal and thereby increasing their shared dynamics.

Notably, due to the subtle differences in clustering patterns across frequencies, we considered microstates C’ and D as identical labels for statistical comparison. Specific labels from the delta band (C’1, C’2), lacking corresponding labels in other bands, were excluded from the mutual information calculation. This exclusion could potentially result in a lower AMI score.

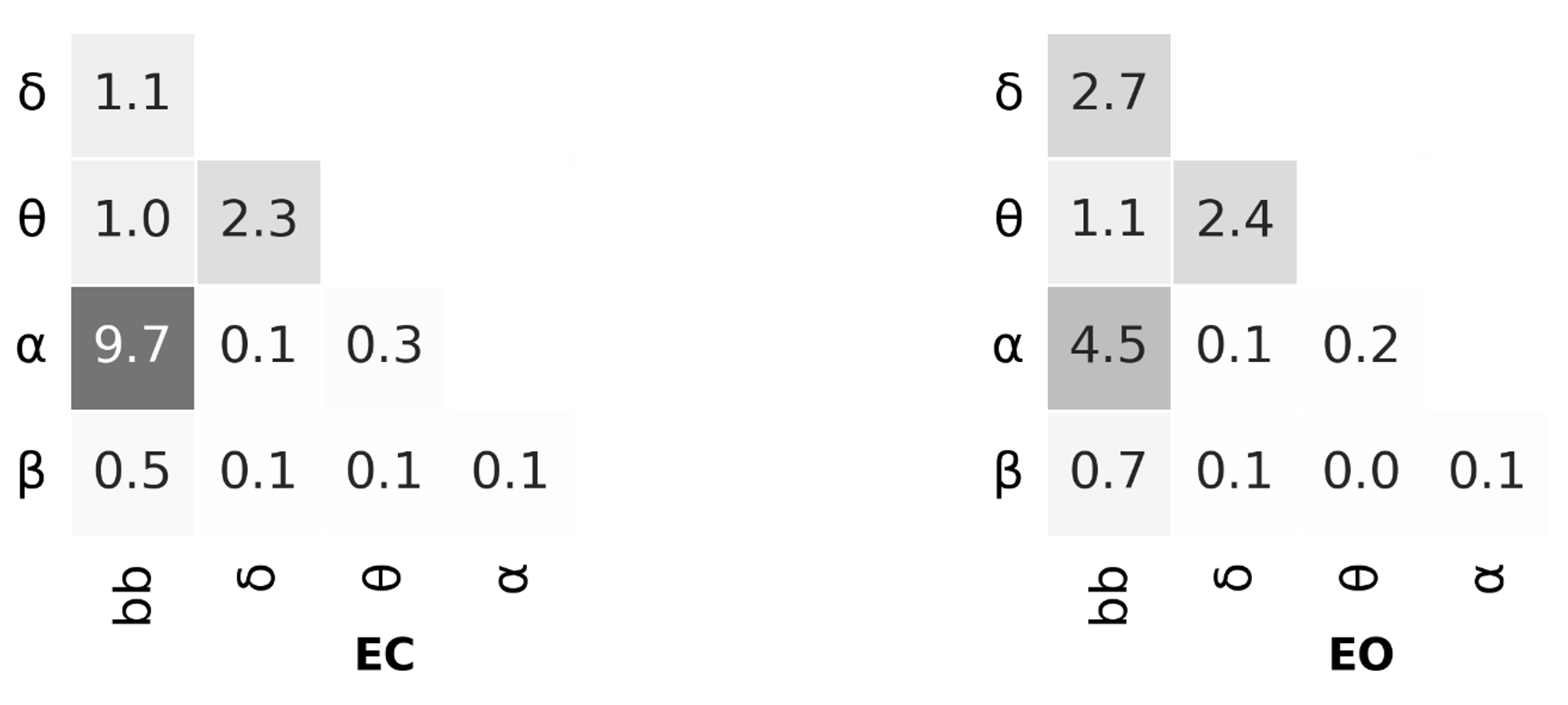

Fig. S6 Mean adjusted mutual information of MS sequences across frequencies (shown in percentage).

Table S1 Statistical description of temporal parameters across various frequency bands under the eyes-closed condition.

|  |  | **delta** | **broad** | **theta** | **alpha** | **beta** |
| --- | --- | --- | --- | --- | --- | --- |
| **A** | GEV | 13.6±2.41 | 9.33±4.13 | 7.99±3.03 | 9.93±5.69 | 9.42±3.11 |
|  | Mean Duration | 0.92±0.04 | 1.05±0.1 | 0.7±0.01 | 1.78±0.28 | 1.17±0.07 |
|  | Time Coverage | 22.89±2.59 | 19.83±6.3 | 15.25±3.58 | 19.68±7.16 | 21.36±4.64 |
|  | Occurrence | 2.24±0.19 | 1.58±0.41 | 2.1±0.47 | 0.92±0.31 | 1.57±0.33 |
| **B** | GEV | 7.67±2.74 | 9.36±4.14 | 17.55±2.57 | 10.57±5.9 | 15.76±3.09 |
|  | Mean Duration | 0.76±0.04 | 1.05±0.11 | 0.66±0.02 | 1.82±0.35 | 1.47±0.12 |
|  | Time Coverage | 11.28±2.6 | 19.36±6.01 | 33.06±2.73 | 20.65±7.41 | 32.0±4.17 |
|  | Occurrence | 1.34±0.25 | 1.54±0.38 | 4.81±0.4 | 0.93±0.3 | 1.83±0.19 |
| **C** | GEV | 25.99±4.65 | 39.16±12.18 | 30.09±5.19 | 44.86±15.38 | 24.06±5.31 |
|  | Mean Duration | 1.04±0.06 | 1.63±0.41 | 0.68±0.02 | 3.14±1.42 | 1.52±0.15 |
|  | Time Coverage | 29.6±3.56 | 49.13±11.14 | 40.73±4.15 | 49.94±13.95 | 38.89±6.32 |
|  | Occurrence | 2.54±0.21 | 2.27±0.32 | 5.71±0.62 | 1.24±0.27 | 2.12±0.26 |
| **D/C'** | GEV | 5.64±1.74 | 4.31±2.62 | 3.74±1.96 | 3.09±2.76 | 2.22±1.49 |
|  | Mean Duration | 0.75±0.03 | 0.92±0.11 | 0.61±0.05 | 1.47±0.23 | 1.09±0.09 |
|  | Time Coverage | 11.01±1.96 | 11.68±5.04 | 10.96±3.6 | 9.72±5.42 | 7.75±3.49 |
|  | Occurrence | 1.32±0.2 | 1.07±0.36 | 1.69±0.53 | 0.55±0.25 | 0.6±0.23 |
| **C'1** | GEV | 7.10 ± 2.29 |  |  |  |  |
|  | Mean Duration | 80.84 ± 5.21 |  |  |  |  |
|  | Time Coverage | 14.46 ± 2.68 |  |  |  |  |
|  | Occurrence | 1.60 ± 0.21 |  |  |  |  |
| **C'2** | GEV | 4.93 ± 1.72 |  |  |  |  |
|  | Mean Duration | 74.78 ± 3.95 |  |  |  |  |
|  | Time Coverage | 10.76 ± 2.24 |  |  |  |  |
|  | Occurrence | 1.30 ± 0.22 |  |  |  |  |

Mean and standard deviation for GEV (in percentage), mean duration (in msec), time coverage (in percentage) occurrence (per second).

Table S2 Statistical description of temporal parameters across various frequency bands under the eyes-open condition.

|  |  | **delta** | **broad** | **theta** | **alpha** | **beta** |
| --- | --- | --- | --- | --- | --- | --- |
| **A** | GEV | 5.72±2.59 | 11.43±4.13 | 8.98±3.24 | 12.64±5.08 | 13.63±3.5 |
|  | Mean Duration | 0.76±0.04 | 1.03±0.09 | 0.7±0.02 | 1.6±0.22 | 1.12±0.08 |
|  | Time Coverage | 10.23±2.36 | 21.88±5.51 | 16.36±3.61 | 22.51±5.6 | 27.8±5.35 |
|  | Occurrence | 1.21±0.23 | 1.8±0.32 | 2.24±0.47 | 1.16±0.23 | 2.15±0.32 |
| **B** | GEV | 10.79±2.45 | 9.8±3.67 | 16.17±3.03 | 13.82±4.89 | 10.97±2.92 |
|  | Mean Duration | 0.93±0.05 | 1.0±0.09 | 0.66±0.02 | 1.65±0.24 | 1.1±0.08 |
|  | Time Coverage | 19.05±2.31 | 19.97±5.68 | 31.34±3.05 | 24.09±5.73 | 23.89±4.53 |
|  | Occurrence | 1.86±0.16 | 1.68±0.34 | 4.53±0.41 | 1.2±0.23 | 1.9±0.28 |
| **C** | GEV | 24.85±4.54 | 31.11±9.26 | 29.77±5.24 | 34.21±11.05 | 27.19±7.4 |
|  | Mean Duration | 1.15±0.08 | 1.42±0.26 | 0.7±0.02 | 2.28±0.67 | 1.13±0.07 |
|  | Time Coverage | 30.44±3.15 | 44.99±9.56 | 39.68±4.07 | 43.15±9.83 | 40.88±8.1 |
|  | Occurrence | 2.37±0.13 | 2.49±0.23 | 5.4±0.48 | 1.52±0.22 | 3.12±0.57 |
| **D/C'** | GEV | 4.43±1.63 | 5.05±2.5 | 4.46±2.01 | 3.43±2.18 | 2.15±1.47 |
|  | Mean Duration | 0.74±0.03 | 0.9±0.07 | 0.62±0.02 | 1.33±0.18 | 0.91±0.06 |
|  | Time Coverage | 8.38±1.9 | 13.17±4.65 | 12.63±3.59 | 10.25±4.02 | 7.43±3.47 |
|  | Occurrence | 1.02±0.19 | 1.27±0.36 | 1.94±0.51 | 0.66±0.21 | 0.72±0.29 |
| **C'1** | GEV | 7.94 ± 2.29 |  |  |  |  |
|  | Mean Duration | 89.25 ± 5.39 |  |  |  |  |
|  | Time Coverage | 16.95 ± 2.76 |  |  |  |  |
|  | Occurrence | 1.72 ± 0.20 |  |  |  |  |
| **C'2** | GEV | 9.85 ± 2.75 |  |  |  |  |
|  | Mean Duration | 86.35 ± 5.57 |  |  |  |  |
|  | Time Coverage | 14.95 ± 2.83 |  |  |  |  |
|  | Occurrence | 1.54 ± 0.20 |  |  |  |  |

Mean and standard deviation for GEV (in percentage), mean duration (in msec), time coverage (in percentage) occurrence (per second).

Table S3 Descriptive statistics: a partial correlation analysis between temporal parameters and scales of emotional disorder (eyes-closed condition).

|  |  | **broadband** | | **delta** | | **theta** | | **alpha** | | **beta** | |
| --- | --- | --- | --- | --- | --- | --- | --- | --- | --- | --- | --- |
|  |  | **Anxiety** | **Depression** | **Anxiety** | **Depression** | **Anxiety** | **Depression** | **Anxiety** | **Depression** | **Anxiety** | **Depression** |
| **A** | GEV | 0.006 | 0.036 | -0.081 | -0.138 | 0.034 | 0.017 | -0.021 | 0.029 | -0.076 | 0.016 |
|  | Mean Duration | 0.071 | 0.007 | -0.048 | -0.144 | 0.002 | 0.037 | 0.109 | -0.002 | -0.034 | 0.012 |
|  | Time Coverage | -0.027 | 0.017 | -0.062 | -0.171 | 0.015 | 0.010 | -0.075 | 0.029 | -0.088 | -0.014 |
|  | Occurrence | -0.087 | 0.009 | -0.054 | -0.145 | 0.015 | 0.006 | -0.145 | 0.010 | -0.068 | -0.020 |
| **B** | GEV | 0.016 | 0.168 | 0.028 | -0.041 | -0.006 | -0.004 | 0.017 | 0.194* | 0.097 | 0.002 |
|  | Mean Duration | 0.087 | 0.134 | 0.080 | -0.056 | -0.021 | 0.033 | 0.14 | 0.133 | 0.124 | 0.012 |
|  | Time Coverage | -0.000 | 0.150 | 0.026 | -0.059 | 0.016 | 0.014 | -0.002 | 0.193* | 0.079 | -0.021 |
|  | Occurrence | -0.064 | 0.098 | 0.019 | -0.045 | 0.020 | 0.002 | -0.110 | 0.100 | -0.011 | -0.037 |
| **C** | GEV | 0.054 | -0.098 | 0.036 | -0.049 | 0.023 | -0.015 | 0.084 | -0.094 | 0.064 | 0.04 |
|  | Mean Duration | 0.118 | -0.102 | 0.056 | -0.015 | -0.045 | 0.072 | 0.179 | -0.102 | 0.127 | 0.011 |
|  | Time Coverage | 0.071 | -0.091 | 0.004 | -0.005 | 0.039 | -0.019 | 0.120 | -0.105 | 0.064 | 0.064 |
|  | Occurrence | -0.133 | 0.064 | -0.027 | 0.005 | 0.050 | -0.038 | -0.155 | 0.047 | -0.029 | 0.076 |
| **D/C'** | GEV | -0.113 | -0.007 | -0.037 | -0.012 | -0.060 | 0.002 | -0.181 | -0.049 | -0.084 | -0.052 |
|  | Mean Duration | -0.071 | -0.023 | 0.016 | -0.032 | 0.023 | -0.010 | -0.059 | -0.041 | -0.020 | 0.002 |
|  | Time Coverage | -0.119 | -0.000 | -0.010 | -0.007 | -0.071 | 0.001 | -0.202* | -0.034 | -0.095 | -0.069 |
|  | Occurrence | -0.125 | 0.009 | -0.013 | 0.002 | -0.070 | 0.000 | -0.221* | -0.025 | -0.095 | -0.082 |
| **C'1** | GEV |  |  | 0.051 | 0.158 |  |  |  |  |  |  |
|  | Mean Duration | |  | 0.078 | 0.137 |  |  |  |  |  |  |
|  | Time Coverage | |  | 0.038 | 0.128 |  |  |  |  |  |  |
|  | Occurrence |  |  | 0.021 | 0.107 |  |  |  |  |  |  |
| **C'2** | GEV |  |  | 0.022 | 0.163 |  |  |  |  |  |  |
|  | Mean Duration | |  | -0.007 | 0.088 |  |  |  |  |  |  |
|  | Time Coverage | |  | 0.000 | 0.127 |  |  |  |  |  |  |
|  | Occurrence |  |  | 0.001 | 0.127 |  |  |  |  |  |  |

Significant correlations (p ≤ .05, FDR control for 32 statistical tests) are denoted by an asterisk (*).

Table S4 Descriptive statistics: a partial correlation analysis between transition probabilities and scales of disorder (eyes-closed condition).

|  |  | **broadband** | | **delta** | | **theta** | | **alpha** | | **beta** | |
| --- | --- | --- | --- | --- | --- | --- | --- | --- | --- | --- | --- |
|  |  | **Anxiety** | **Depression** | **Anxiety** | **Depression** | **Anxiety** | **Depression** | **Anxiety** | **Depression** | **Anxiety** | **Depression** |
| **From A To:** | B | 0.017 | 0.113 | 0.022 | -0.065 | 0.017 | 0.035 | -0.034 | 0.15 | 0.136 | -0.014 |
|  | C | 0.056 | -0.096 | -0.033 | -0.027 | -0.03 | 0.065 | 0.126 | -0.101 | 0.048 | 0.133 |
|  | D/C' | -0.085 | -0.004 | -0.015 | -0.016 | -0.133 | -0.027 | -0.154 | -0.044 | -0.113 | -0.089 |
|  | C'1 |  |  | 0.012 | 0.083 |  |  |  |  |  |  |
|  | C'2 |  |  | -0.003 | 0.114 |  |  |  |  |  |  |
| **From B To:** | A | -0.023 | 0.016 | -0.05 | -0.166 | 0.044 | 0.098 | -0.083 | -0.03 | 0.01 | 0.033 |
|  | C | 0.094 | 0.001 | -0.018 | -0.01 | 0.04 | -0.043 | 0.198 | 0.026 | 0.107 | 0.059 |
|  | D/C' | -0.098 | -0.024 | -0.007 | -0.004 | -0.091 | -0.016 | -0.218* | -0.01 | -0.073 | -0.075 |
|  | C'1 |  |  | 0.024 | 0.097 |  |  |  |  |  |  |
|  | C'2 |  |  | 0.006 | 0.124 |  |  |  |  |  |  |
| **From C To:** | A | 0.011 | -0.097 | -0.058 | -0.142 | -0.04 | 0.068 | 0.028 | -0.059 | -0.022 | 0.014 |
|  | B | 0.059 | 0.1 | 0.026 | -0.049 | -0.006 | -0.013 | 0.148 | 0.105 | 0.106 | 0.024 |
|  | D/C' | -0.043 | -0.007 | -0.011 | 0 | -0.06 | -0.016 | -0.17 | -0.055 | -0.072 | -0.063 |
|  | C'1 |  |  | 0.015 | 0.095 |  |  |  |  |  |  |
|  | C'2 |  |  | 0 | 0.128 |  |  |  |  |  |  |
| **From D/C' To:** | A | -0.073 | -0.04 | -0.062 | -0.171 | -0.057 | -0.011 | -0.06 | -0.073 | 0.007 | 0.007 |
|  | B | -0.002 | 0.085 | 0.028 | -0.052 | 0.048 | 0.03 | -0.024 | 0.094 | 0.013 | 0.009 |
|  | C | 0.044 | -0.027 | -0.027 | 0 | -0.018 | -0.035 | 0.07 | -0.032 | 0.116 | 0.123 |
|  | C'1 |  |  | 0.013 | 0.095 |  |  |  |  |  |  |
|  | C'2 |  |  | -0.001 | 0.119 |  |  |  |  |  |  |
| **From C'1 To:** | A |  |  | -0.063 | -0.162 |  |  |  |  |  |  |
|  | B |  |  | 0.035 | -0.04 |  |  |  |  |  |  |
|  | C |  |  | -0.021 | 0.023 |  |  |  |  |  |  |
|  | D/C' |  |  | -0.01 | 0.014 |  |  |  |  |  |  |
|  | C'2 |  |  | 0.001 | 0.128 |  |  |  |  |  |  |
| **From C'2 To:** | A |  |  | -0.067 | -0.16 |  |  |  |  |  |  |
|  | B |  |  | 0.03 | -0.037 |  |  |  |  |  |  |
|  | C |  |  | -0.027 | 0.03 |  |  |  |  |  |  |
|  | D/C' |  |  | -0.011 | 0.016 |  |  |  |  |  |  |
|  | C'1 |  |  | 0.014 | 0.105 |  |  |  |  |  |  |

Significant correlations (p ≤ .05, FDR control for 24 statistical tests) are denoted by an asterisk (*).

**4 Age and gender effects on microstate parameters**

**Age difference.** We applied independent sample comparisons to investigate differences in age (younger vs older) and gender (male vs female) groups, results are depicted in Fig. S7.

In broadband, all parameters (GEV, duration, coverage, occurrence) were observed to increase as a function of age in microstates A and B, while decrease in microstate C. A similar trend was also found in narrowband, with an exception of theta. Older individuals showed minor differences in the theta band compared to younger adults, with no significant variations detected during EO conditions. Additionally, certain effects in the narrowband were statistically equivalent to zero, which was not observed in the broadband.

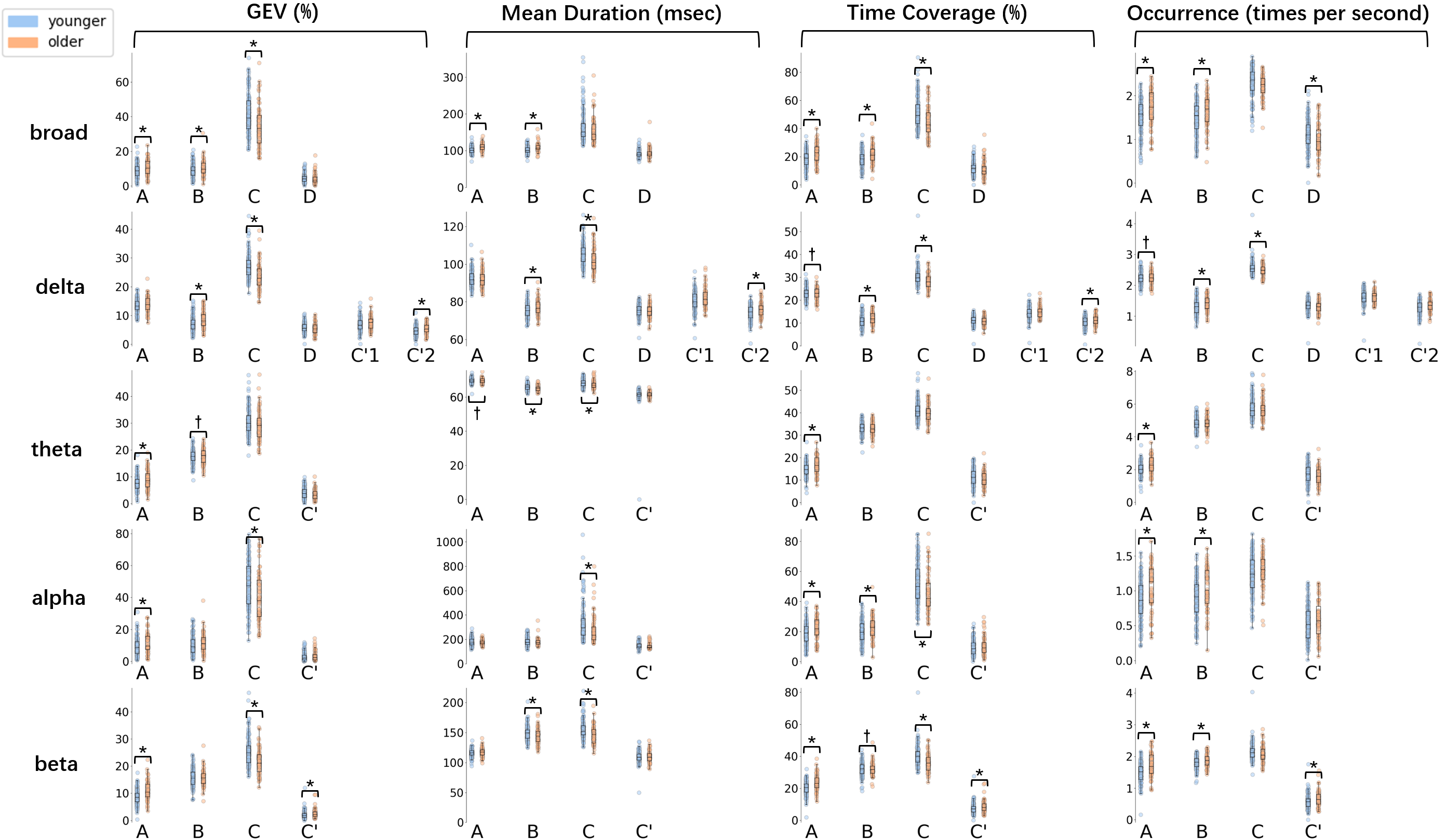

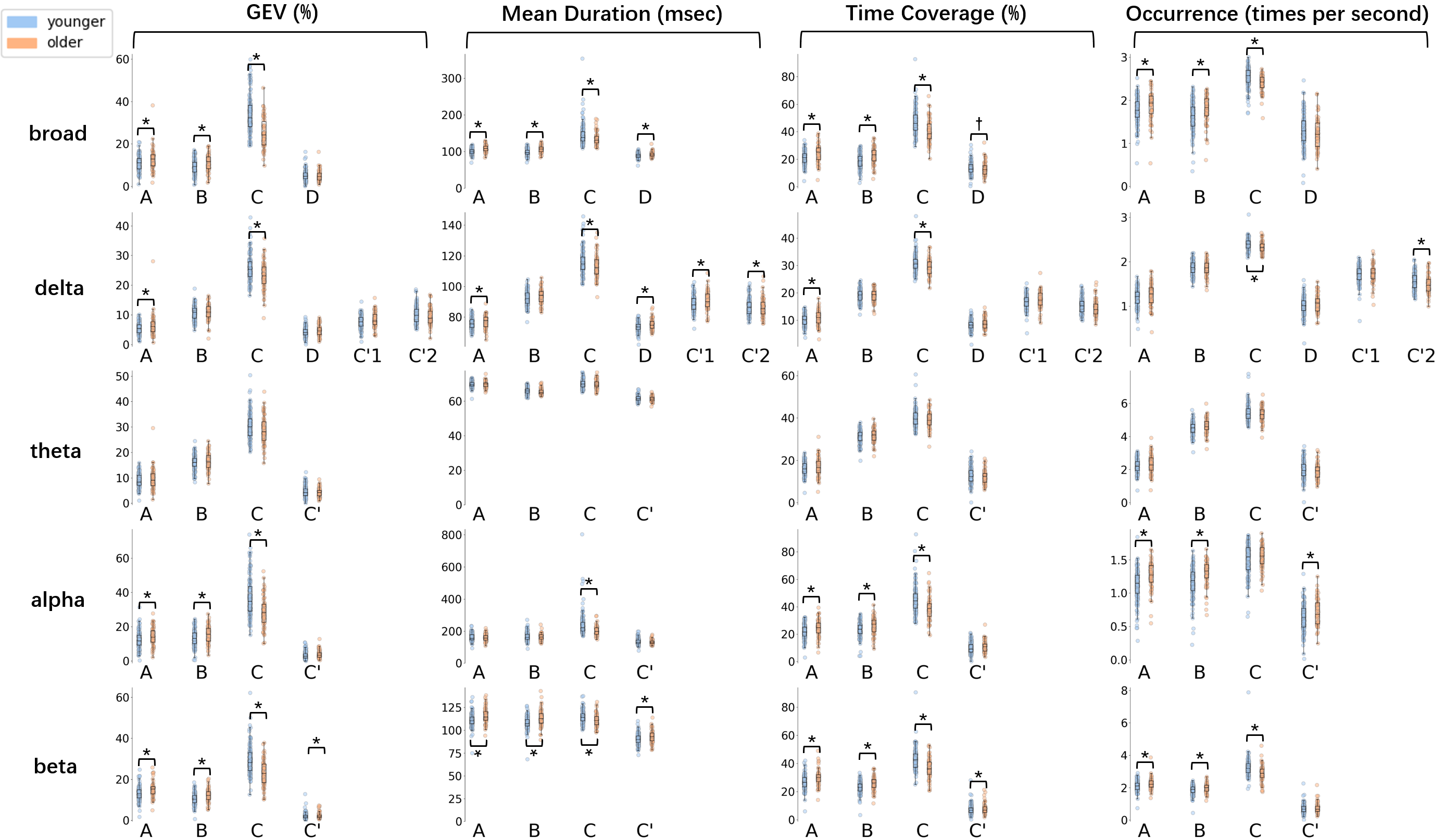

Fig. S7 Comparison between younger and older individuals across all frequency bands (top: EC condition; bottom: EO condition). Significance was denoted by "*", corresponding to corrected p ≤ 0.05. Parameters with no significant difference were marked with "†". Standardized Mean Difference was used to represent the magnitude of differences.

**Gender difference.** No significant differences were observed between male and female, with the exception of the delta band: Compared to males, females tended to exhibit higher values in microstates C’1 and C’2, while lower in microstates A and B. Notably, microstate D in the delta frequency band showed statistical equivalence to zero (d < .05). These findings reveal potential gender-specific predictors in narrowband EEG. Fig. S8 plots the individual participant data for gender comparisons.

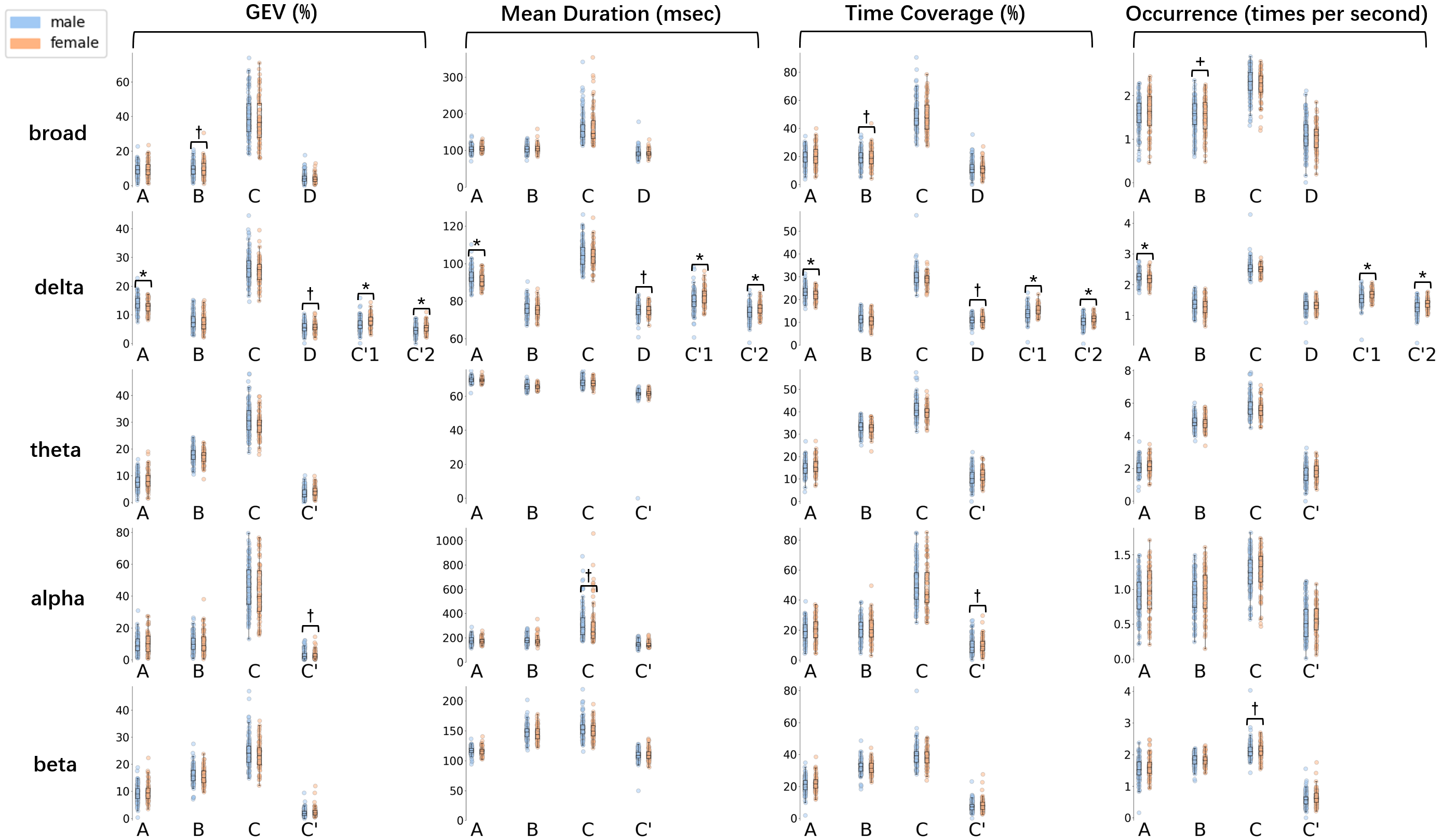

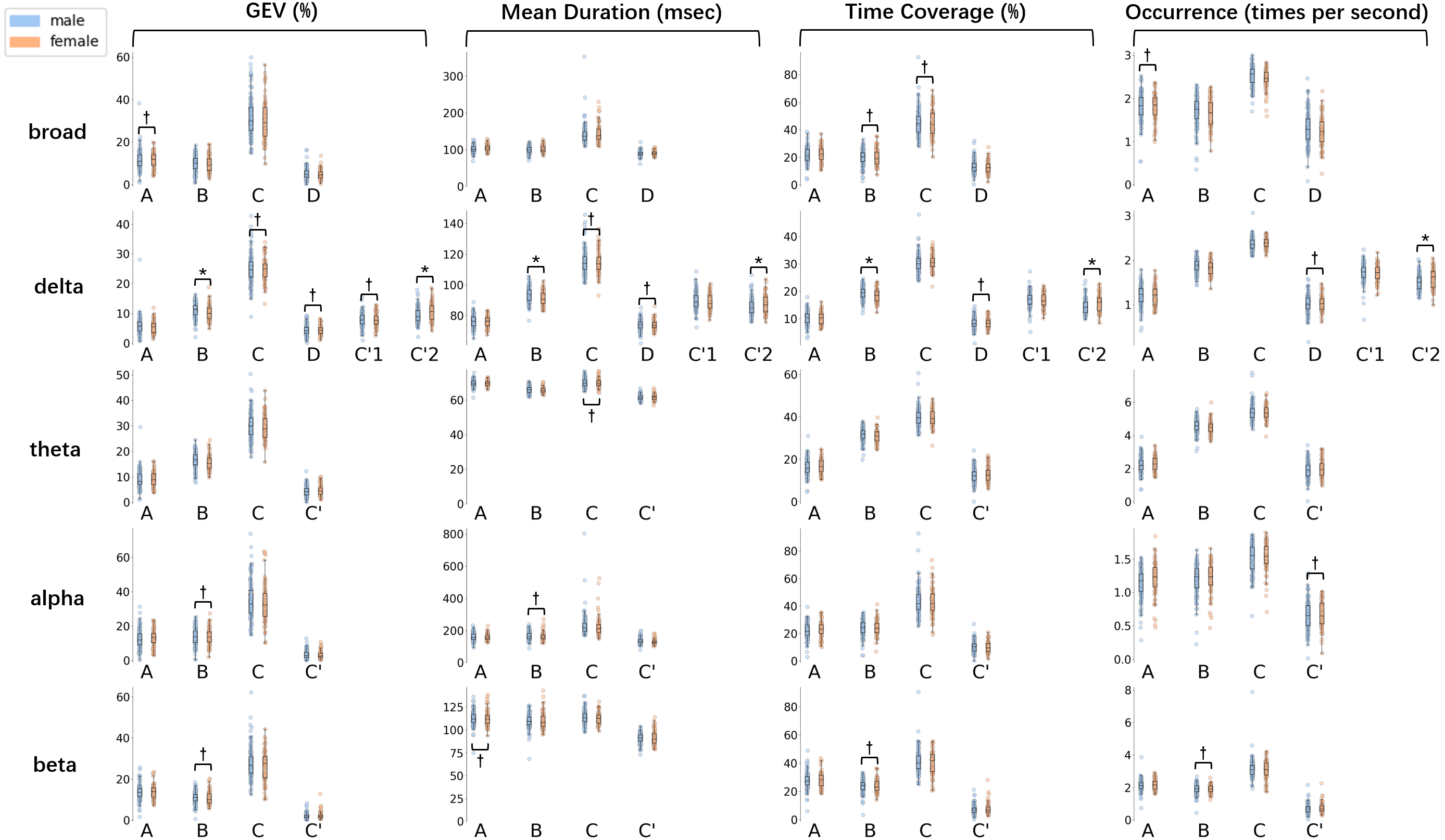

Fig. S8 Comparison between males and females across all frequency bands (top: EC condition; bottom: EO condition). Significance was denoted by "*", corresponding to corrected p ≤ 0.05. Parameters with no significant difference were marked with "†". Standardized Mean Difference was used to represent the magnitude of differences.

**Age and gender effect on microstate parameters.** In this study, we observed a general tendency for increased parameters as a function of age, except for microstate C. This pattern aligns with (Tomescu et al., 2018), who analyzed eyes-closed EEG recordings from a wider age range (6–87 years). However, our present finding of the increased occurrence in microstates A and B was inconsistent, as they reported a total decrease. (Zanesco et al., 2020) also reported higher GEV in microstates A and B and lower in microstate C among older individuals. However, they observed fewer occurrences with age, while we found increased occurrences in microstates A and B. These differences could be attributed to the determination of the resting state. As they analyzed average parameters across EC and EO, we conducted comparisons for each condition separately. Another potential explanation could be the different number of clusters.

We also examined the gender effect on microstates, revealing a weak association between them. Gender differences were only apparent in the delta band, where females tend to exhibit higher values in microstates C’1 and C’2, but lower in microstates A and B. In the broadband, we observed a general trend for females to exhibit longer durations. This finding partially aligns with (Tomescu et al., 2018) and (Zanesco et al., 2020), who analyzed a similar cohort and reported more prolonged durations in females. In contrast, the absence of significance in our study might be due to methodological differences, as we calculated additional temporal parameters and applied FDR correction.

Overall, these results appear to support the ideas of Koenig and colleagues, who indicated that microstate profiles follow a lawful, but complex behavior with age and gender (Koenig et al., 2002). Further research with broader developmental cohorts is needed to explore this effect.

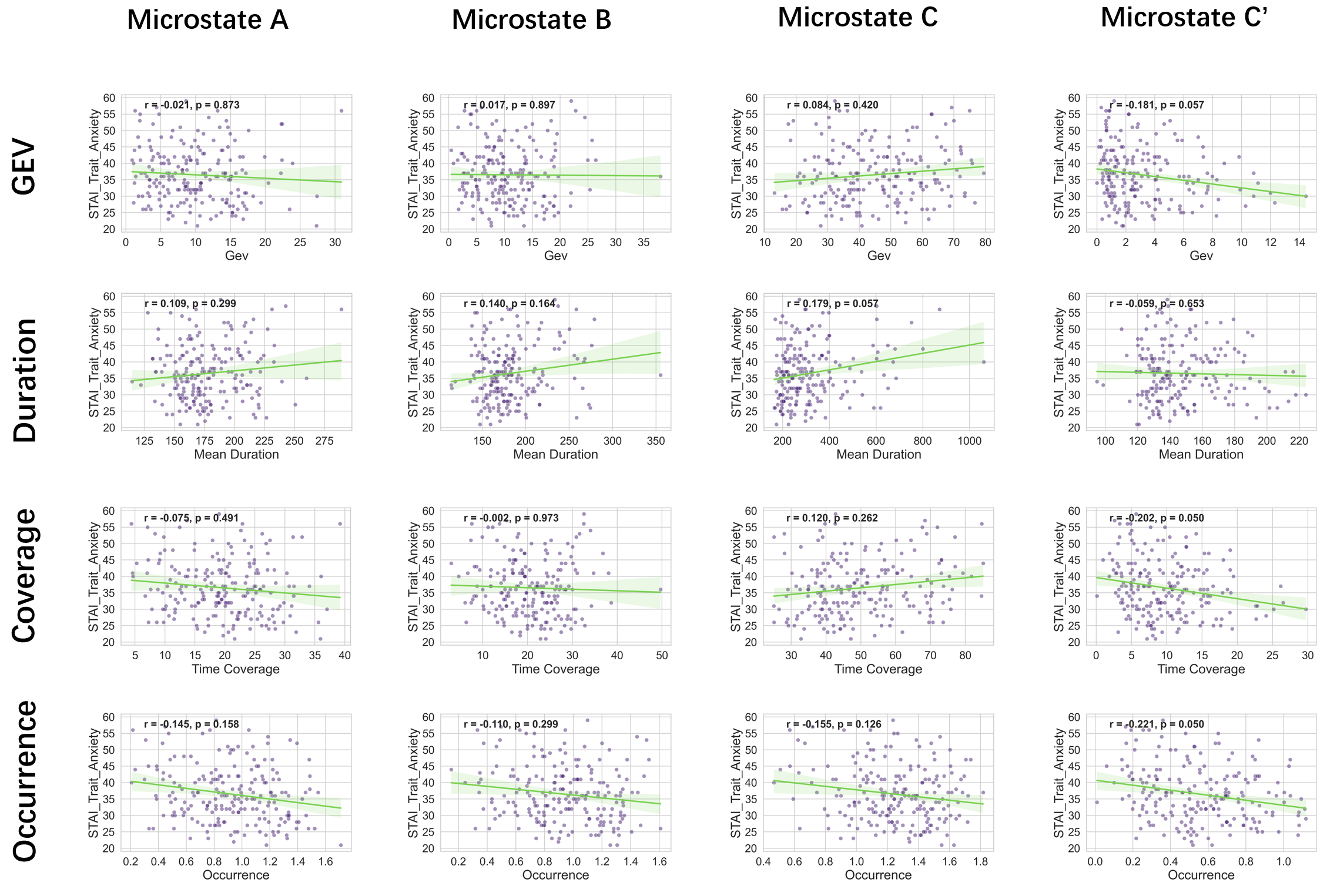

Fig. S9 Plots of the correlation analysis between trait-anxiety and each of the four microstate parameters during eyes-closed condition (alpha band, FDR corrected).

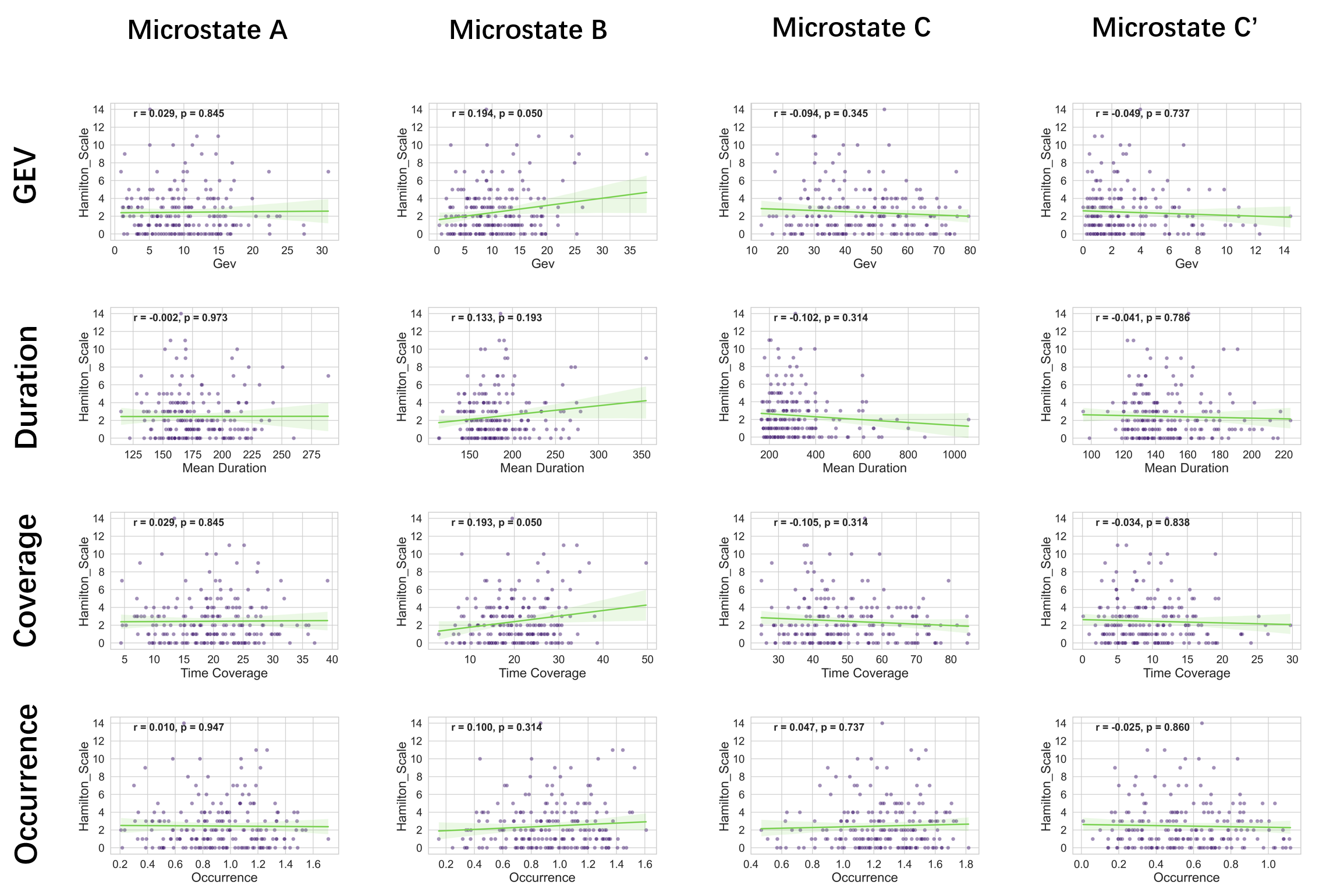

Fig. S10 Plots of the correlation analysis between Hamilton Scales and each of the four microstate parameters during eyes-closed condition (alpha band, FDR corrected).

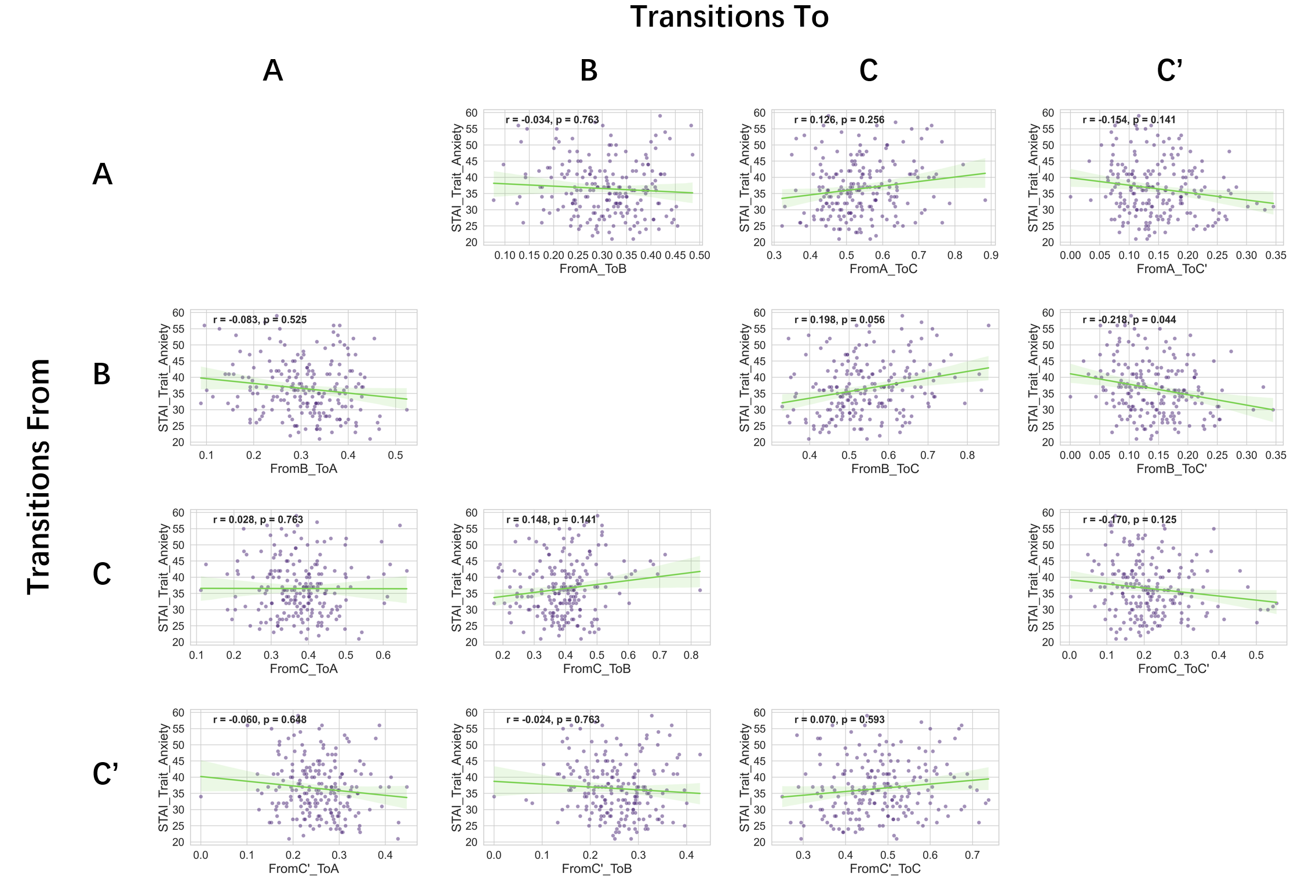

Fig. S11 Plots of the correlation analysis between trait-anxiety and each of the transition probability during eyes-closed condition (alpha band, FDR corrected).

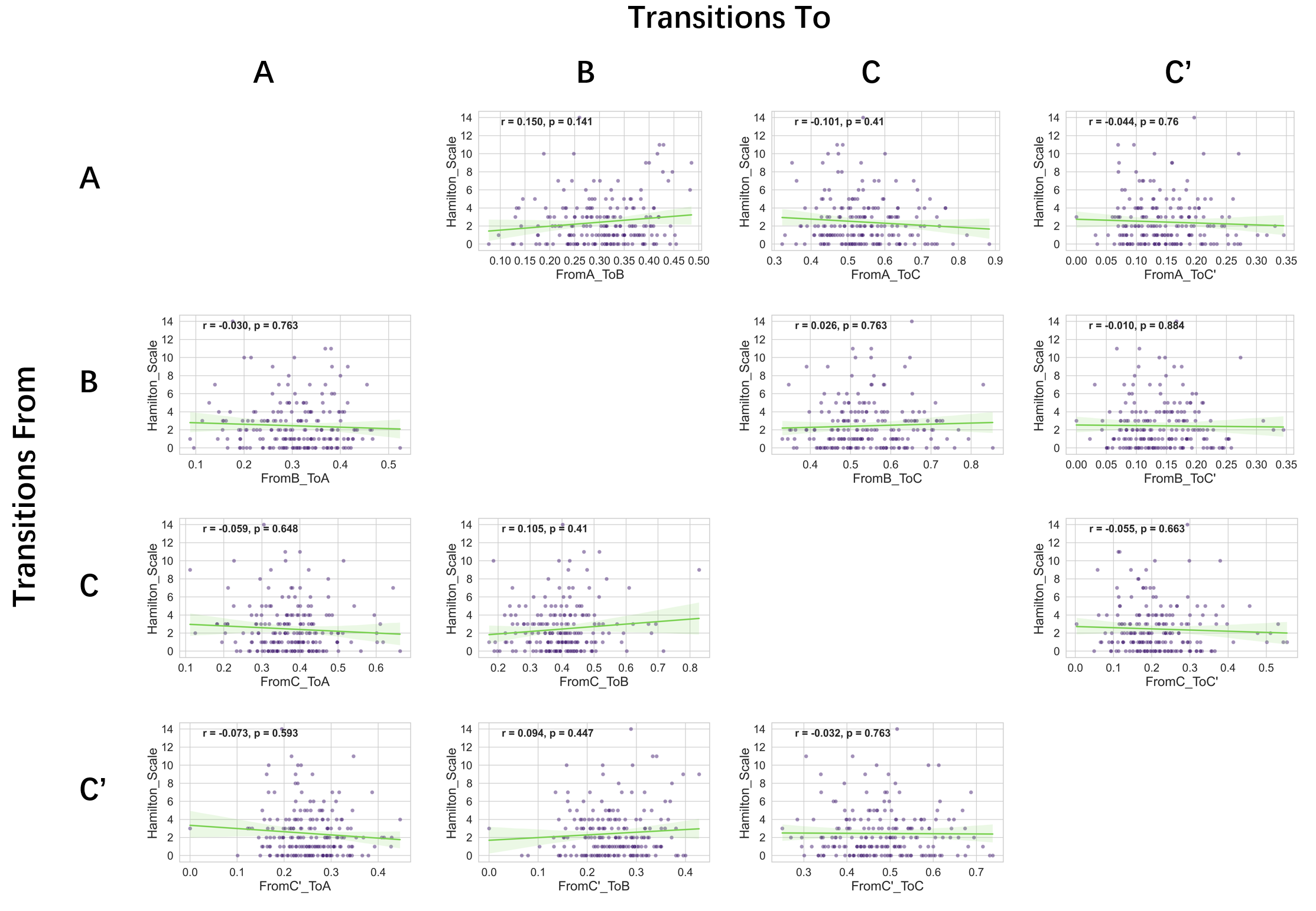

Fig. S12 Plots of the correlation analysis between Hamilton Scales and each of the transition probability during eyes-closed condition (alpha band, FDR corrected).

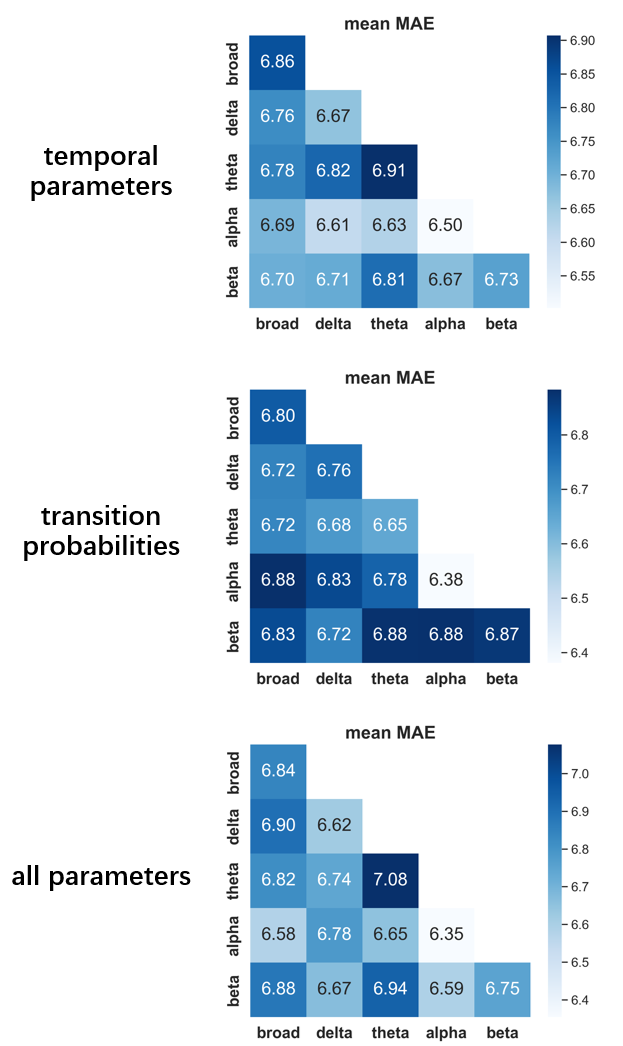

Fig. S13 Performance of the trait anxiety prediction model (eyes-closed condition). Each grid shows the mean MAE of microstate features in one frequency band (on the diagonal) or of the combination of microstate features in two frequency bands (off the diagonal). The mean MAE derived from a five-fold cross-validation procedure (top row: models containing four temporal parameters; middle row: models containing only transition probabilities; bottom row: models containing all microstate parameters).

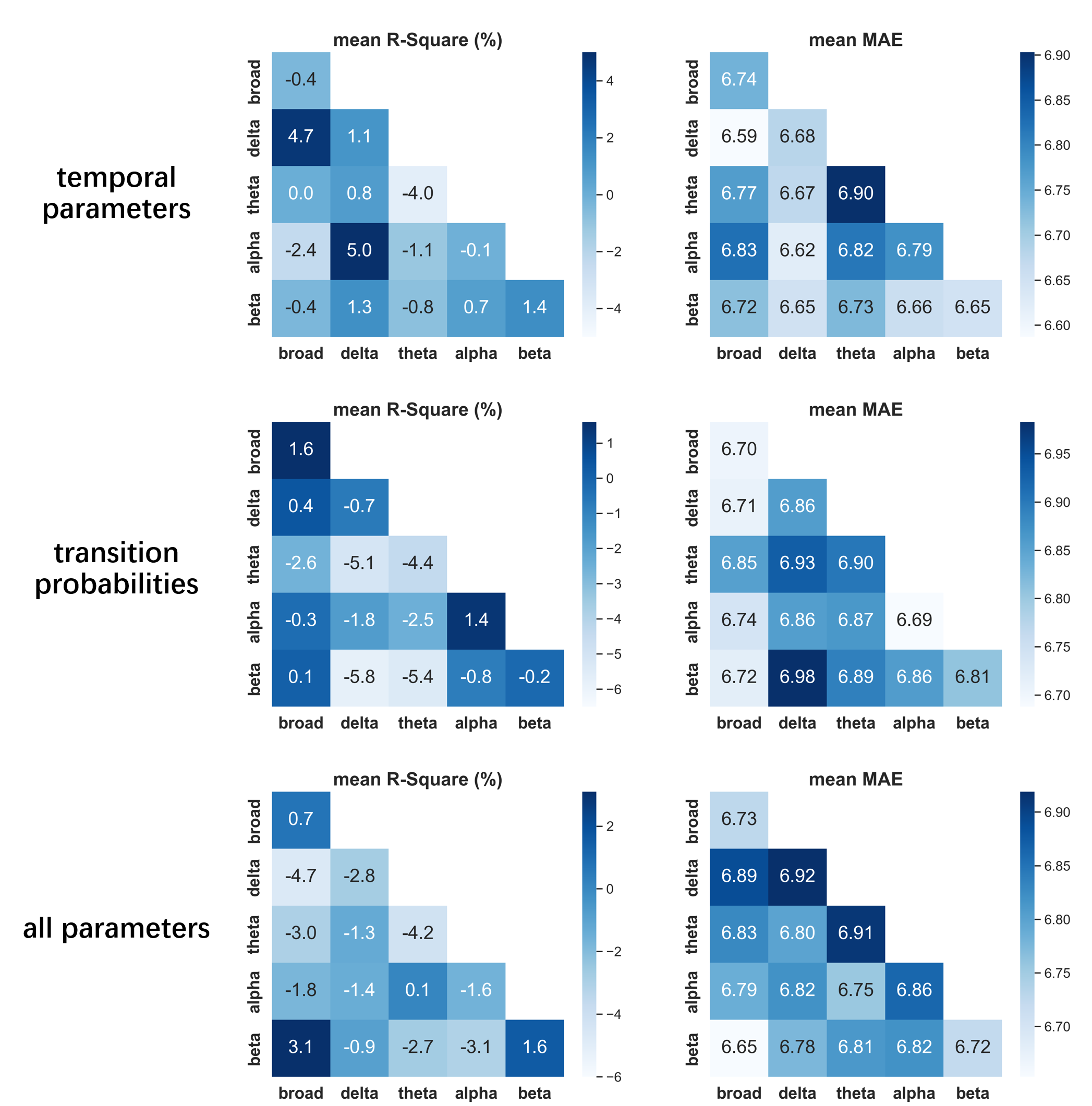

Fig. S14 Performance of the trait anxiety prediction model (eyes-open condition). Each grid shows the mean R-square or MAE of microstate features in one frequency band (on the diagonal) or of the combination of microstate features in two frequency bands (off the diagonal). The mean R-square and MAE derived from a five-fold cross-validation procedure (top row: models containing four temporal parameters; middle row: models containing only transition probabilities; bottom row: models containing all microstate parameters).

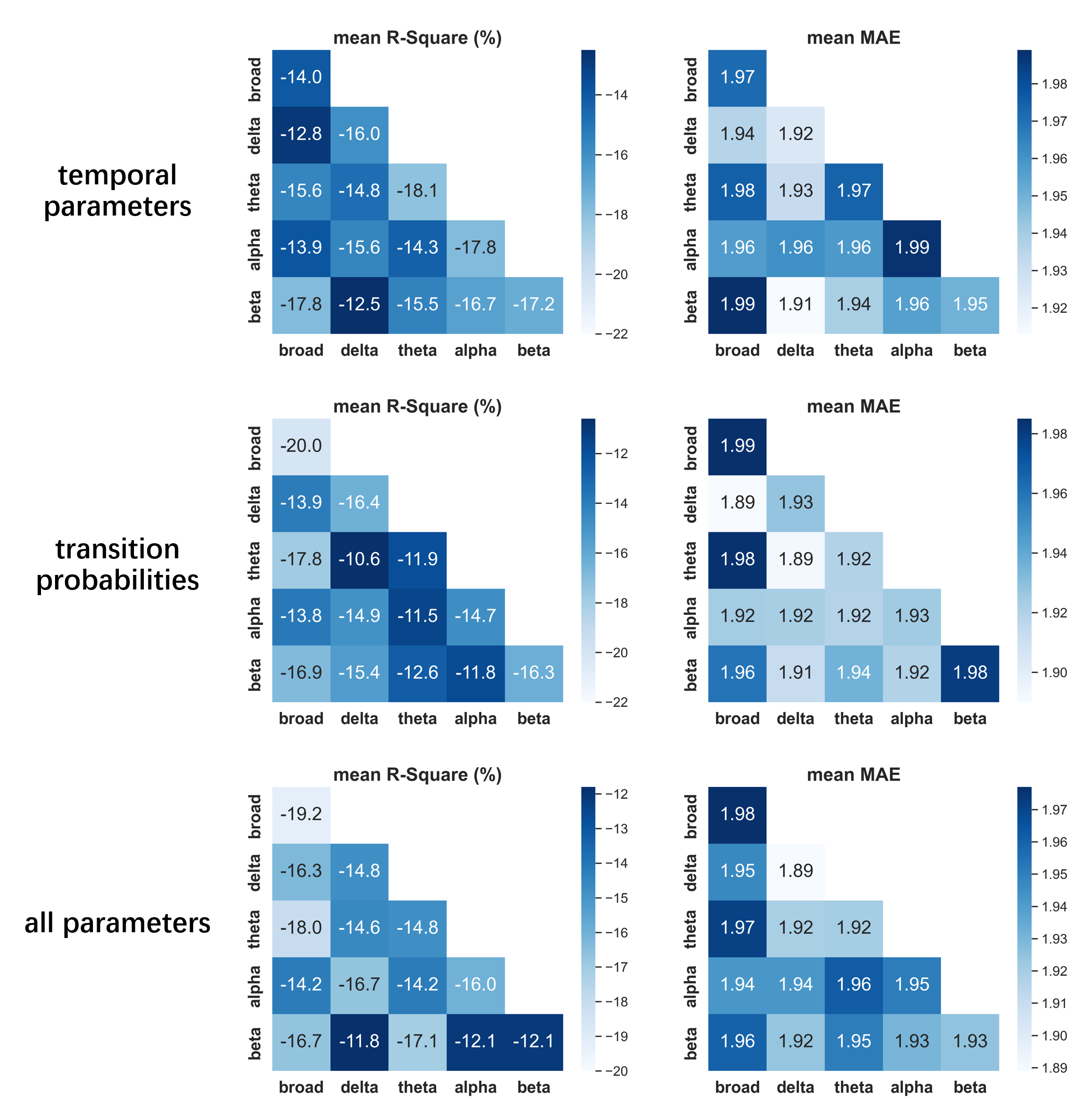

Fig. S15 Performance of the Hamilton scale prediction model (eyes-closed condition). Each grid shows the mean R-square or MAE of microstate features in one frequency band (on the diagonal) or of the combination of microstate features in two frequency bands (off the diagonal). The mean R-square and MAE derived from a five-fold cross-validation procedure (top row: models containing four temporal parameters; middle row: models containing only transition probabilities; bottom row: models containing all microstate parameters).

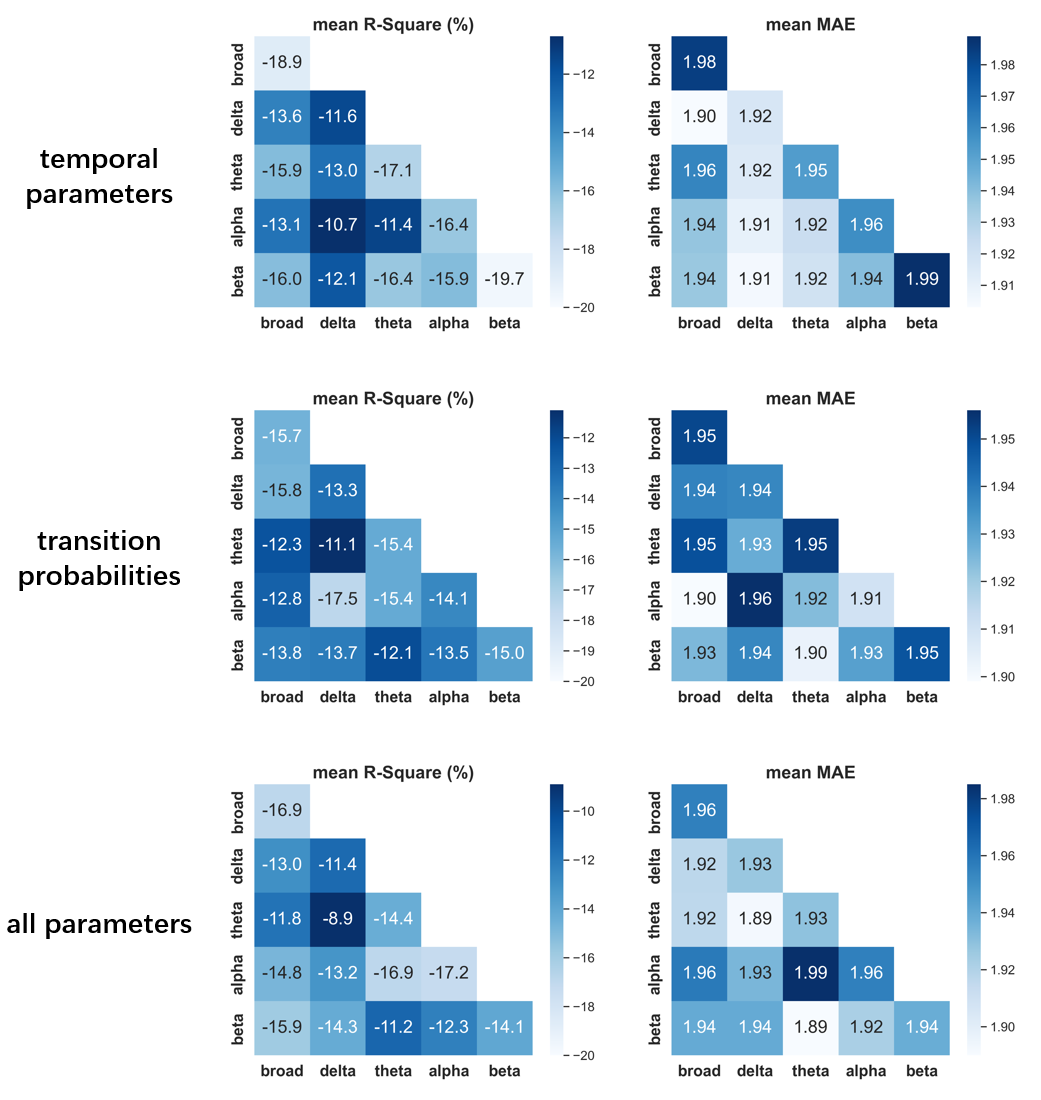

Fig. S16 Performance of the Hamilton scale prediction model (eyes-open condition). Each grid shows the mean R-square or MAE of microstate features in one frequency band (on the diagonal) or of the combination of microstate features in two frequency bands (off the diagonal). The mean R-square and MAE derived from a five-fold cross-validation procedure (top row: models containing four temporal parameters; middle row: models containing only transition probabilities; bottom row: models containing all microstate parameters).

**References**

Custo, A., Van De Ville, D., Wells, W.M., Tomescu, M.I., Brunet, D., Michel, C.M., 2017. Electroencephalographic Resting-State Networks: Source Localization of Microstates. Brain Connect. 7, 671–682. https://doi.org/10.1089/brain.2016.0476

Custo, A., Vulliemoz, S., Grouiller, F., Van De Ville, D., Michel, C., 2014. EEG source imaging of brain states using spatiotemporal regression. Neuroimage 96, 106–116. https://doi.org/10.1016/j.neuroimage.2014.04.002

Dale, A.M., Fischl, B., Sereno, M.I., 1999. Cortical Surface-Based Analysis: I. Segmentation and Surface Reconstruction. NeuroImage 9, 179–194. https://doi.org/10.1006/nimg.1998.0395

Goldman, R., Cohen, M., Stern, J., Engel, J., 2001. Tomographic Mapping of Alpha Rhythm Using Simultaneous EEG/fMRI. Presented at the Neuroimage. https://doi.org/10.1016/S1053-8119(01)92605-9

Groppe, D.M., Bickel, S., Keller, C.J., Jain, S.K., Hwang, S.T., Harden, C., Mehta, A.D., 2013. Dominant frequencies of resting human brain activity as measured by the electrocorticogram. NeuroImage 79, 223–233. https://doi.org/10.1016/j.neuroimage.2013.04.044

Helmholtz, H., 1853. Ueber einige Gesetze der Vertheilung elektrischer Ströme in körperlichen Leitern mit Anwendung auf die thierisch-elektrischen Versuche. Annalen der Physik 165, 211–233. https://doi.org/10.1002/andp.18531650603

Koenig, T., Prichep, L., Lehmann, D., Sosa, P.V., Braeker, E., Kleinlogel, H., Isenhart, R., John, E.R., 2002. Millisecond by Millisecond, Year by Year: Normative EEG Microstates and Developmental Stages. NeuroImage 16, 41–48. https://doi.org/10.1006/nimg.2002.1070

Mantini, D., Perrucci, M.G., Del Gratta, C., Romani, G.L., Corbetta, M., 2007. Electrophysiological signatures of resting state networks in the human brain. Proc. Natl. Acad. Sci. U. S. A. 104, 13170–13175. https://doi.org/10.1073/pnas.0700668104

Mellem, M.S., Wohltjen, S., Gotts, S.J., Ghuman, A.S., Martin, A., 2017. Intrinsic frequency biases and profiles across human cortex. Journal of Neurophysiology 118, 2853–2864. https://doi.org/10.1152/jn.00061.2017

Michel, C.M., Murray, M.M., Lantz, G., Gonzalez, S., Spinelli, L., Grave de Peralta, R., 2004. EEG source imaging. Clinical Neurophysiology 115, 2195–2222. https://doi.org/10.1016/j.clinph.2004.06.001

Tadel, F., Baillet, S., Mosher, J.C., Pantazis, D., Leahy, R.M., 2011. Brainstorm: A User-Friendly Application for MEG/EEG Analysis. Computational Intelligence and Neuroscience 2011, e879716. https://doi.org/10.1155/2011/879716

Tomescu, M.I., Rihs, T.A., Rochas, V., Hardmeier, M., Britz, J., Allali, G., Fuhr, P., Eliez, S., Michel, C.M., 2018. From swing to cane: Sex differences of EEG resting-state temporal patterns during maturation and aging. Developmental Cognitive Neuroscience 31, 58–66. https://doi.org/10.1016/j.dcn.2018.04.011

Trevor Hastie, Robert Tibshirani, Jerome Friedman, 2009. The Elements of Statistical Learning.

Zanesco, A.P., King, B.G., Skwara, A.C., Saron, C.D., 2020. Within and between-person correlates of the temporal dynamics of resting EEG microstates. NeuroImage 211, 116631. https://doi.org/10.1016/j.neuroimage.2020.116631
